# Drug and Single-Cell Gene Expression Integration Identifies Sensitive and Resistant Glioblastoma Cell Populations

**DOI:** 10.1101/2025.05.15.654044

**Authors:** Robert K. Suter, Anna M. Jermakowicz, Rithvik Veeramachaneni, Matthew D’Antuono, Longwei Zhang, Rishika Chowdary, Simon Kaeppeli, Madison Sharp, Pravallika Palwai, Vasileios Stathias, Grace Baker, Luz Ruiz, Winston Walters, Maria Cepero, Danielle Burgenske, Edward B Reilly, Anatol Oleksijew, Mark G. Anderson, Sion Ll. Williams, Michael E. Ivan, Ricardo J. Komotar, Macarena I. De La Fuente, Gregory Stein, Alexander Wojcinski, Santosh Kesari, Jann N. Sarkaria, Stephan C. Schürer, Nagi G. Ayad

## Abstract

Glioblastoma (GBM) remains the most common and lethal adult malignant primary brain cancer with few treatment options. A significant issue hindering GBM therapeutic development is intratumor heterogeneity. GBM tumors contain neoplastic cells within a spectrum of different transcriptional states. Identifying effective therapeutics requires a platform that predicts the differential sensitivity and resistance of these states to various treatments. Here, we developed a novel framework, ISOSCELES (<u>I</u>nferred cell <u>S</u>ensitivity <u>O</u>perating on the integration of <u>S</u>ingle-<u>C</u>ell <u>E</u>xpression and <u>L</u>1000 <u>E</u>xpression <u>S</u>ignatures), to quantify the cellular drug sensitivity and resistance landscape. Using single-cell RNA sequencing of newly diagnosed and recurrent GBM tumors, we identified compounds from the LINCS L1000 database with transcriptional response signatures selectively discordant with distinct GBM cell states. We validated the significance of these findings *in vitro, ex vivo,* and *in vivo*, and identified a novel combination of an OLIG2 inhibitor and Depatux-M for GBM. Our studies suggest that ISOSCELES identifies cell states sensitive and resistant to targeted therapies in GBM and that it can be applied to identify new synergistic combinations.

**Highlights:**

- Integration of GBM single-cell RNA sequencing data with L1000-derived drug response signatures facilitates clustering of tumor cells and small molecules on cell-drug connectivity.
- Cell-drug connectivity predicts the identities of drug-sensitive and resistant cell states.
- In silico perturbation analysis using cell-drug connectivity predicts drug-induced changes in the cell-drug connectivity landscape in vivo.
- In silico perturbation analysis to predict drug-induced changes in the tumor cell-drug connectivity landscape predicts drug combinations that synergize in vivo to extend survival.

## Introduction

Glioblastoma (GBM) remains the most common adult malignant brain cancer, with a median overall survival of only 15 months.^1–3^ No new targeted therapies have been approved for GBM since the introduction of the alkylating agent temozolomide (TMZ) in 2005.^1–3^ Thus, novel therapeutic options are direly needed for patients with GBM.

Intratumor heterogeneity poses a significant barrier to identifying effective targeted therapies for GBM.^4,5^ Single GBM tumors contain diverse populations of cells with considerable genomic, transcriptomic, and proteomic differences.^6–9^ GBM single-cell characterization has revealed a plastic transcriptional spectrum reminiscent of canonical neurodevelopmental cell types.^6^ These transcriptional states encompass astrocyte-like (AC-like), neural-progenitor-like (NPC-like), oligodendrocyte-progenitor-like (OPC-like), and mesenchymal-like (MES-like) cells. The importance of their relative abundance is highlighted by the identification of transcriptional subtype (Classical, Proneural, Mesenchymal) plasticity and switching following standard-of-care treatment.^10^ Therefore, it is necessary to develop novel means of targeting these dynamic tumor cell populations. However, there is no current method to predict which compounds target the different neoplastic cell states within GBM tumors. We and others have shown that cancer gene expression-based disease signature reversal is predictive of compound efficacy^11–13^. Using the NIH Library for Integrated Network-based Cellular Signatures (LINCS) L1000 assay dataset, we previously developed transcriptional consensus signatures (TCSs) for each small molecule, consisting of genes that are consistently up- or down-regulated irrespective of cell type. We showed that effective combination treatments can be identified through the integration of compound TCSs with TCGA-derived GBM disease signatures and that compound efficacy in a GBM subtype-specific manner can be predicted.^12–14^ However, these analyses do not address the GBM intratumor heterogeneity that is evident after performing single-cell RNA sequencing (scRNAseq), to predict mechanisms of drug sensitivity or resistance.

Here, we posit a novel framework for the integration of scRNAseq data and L1000 TCSs to facilitate *in silico* perturbation analyses, ISOSCELES (<u>I</u>nferred cell <u>S</u>ensitivity <u>O</u>perating on the integration of <u>S</u>ingle-<u>C</u>ell <u>E</u>xpression and <u>L</u>1000 <u>E</u>xpression <u>S</u>ignatures). Using ISOSCELES, we demonstrate that the integration of GBM single-cell disease signatures with LINCS L1000-derived compound response signatures can identify drugs most likely to target transcriptionally distinct cell populations in individual GBM tumors. Furthermore, we demonstrate that this framework can predict the sensitive and resistant cell populations within GBM tumors in an orthotopic xenograft model. Using patient single-cell RNA sequencing data and a TCS for the aurora kinase inhibitor alisertib, we predict and confirm both the MES-like transcriptional identity of a distinct alisertib-resistant cell population and the depletion of transcriptionally NPC-like cells *in vivo.*^15^ Moreover, we use publicly available datasets to demonstrate the power of this approach to predict treatment-induced transcriptional response in discrete GBM cell populations and to identify synergistic small molecule combinations. Building on this, we leverage our framework to identify a combination of a novel OLIG2 inhibitor, CT-179, with an anti-EGFR antibody-drug conjugate, Depatuxizumab Mafodotin (Depatux-M, ABT-414), which synergizes to increase survival in an orthotopic xenograft model of GBM. Importantly, we introduce ISOSCELES as a publicly available R package and Shiny application. ISOSCELES uses the methods defined here to generate and analyze cell-type-specific disease signatures, perform *in silico* drug connectivity analyses, to characterize cell population sensitivity or resistance to specific treatments, and to prioritize synergistic combinations.

## Results

To identify the different cell types within GBM tumors, we performed single-cell RNA-sequencing of six samples resected from three newly diagnosed and three recurrent GBM patients. We combined our sequencing results with those from published studies that had used the same 10X Genomics platform to study GBM at the single cell level to increase sample size for our downstream analyses **(Fig. 1)**.^16^ Our dataset (hereto referred to as Suter et al.) and the Johnson et al. dataset were integrated to reveal a large tumor cell population **(Fig. 1d)**.^17^ This population contained tumor cells spanning the neurodevelopmentally-rooted GBM cell transcriptional states identified by Neftel et al **(Fig. 1e-f)**.^6^ In addition to neoplastic cells, both datasets contain populations of myeloid cells, T-cells, fibroblasts, endothelial cells, and oligodendrocytes. These non-neoplastic cell populations that comprise the tumor microenvironment (TME) were identified by the expression of specific marker genes, such as PTPRC for myeloid cells and CD3E for T-cells **(Supplementary Fig. S1)**, and contain cells captured across both datasets **(Fig. 1g)**. We then designated the transcriptional profiles of these TME cells as our “normal” cell reference to be able to compare the distinct GBM tumor cell states represented within our dataset. Utilizing this comparison, we were able to derive a pseudo-bulk disease signature for each of the cell states within GBM tumors **(Supplementary Fig. S1h)**. The disease signatures unique to each transcriptional state reflected previously reported expression markers, such as CDK4 and SOX4 overexpression in NPC- and OPC-like cells, EGFR overexpression in AC-like cells, and CD44 overexpression in MES-like cells **(Supplementary Fig. S1h)**.

**Figure 1:**
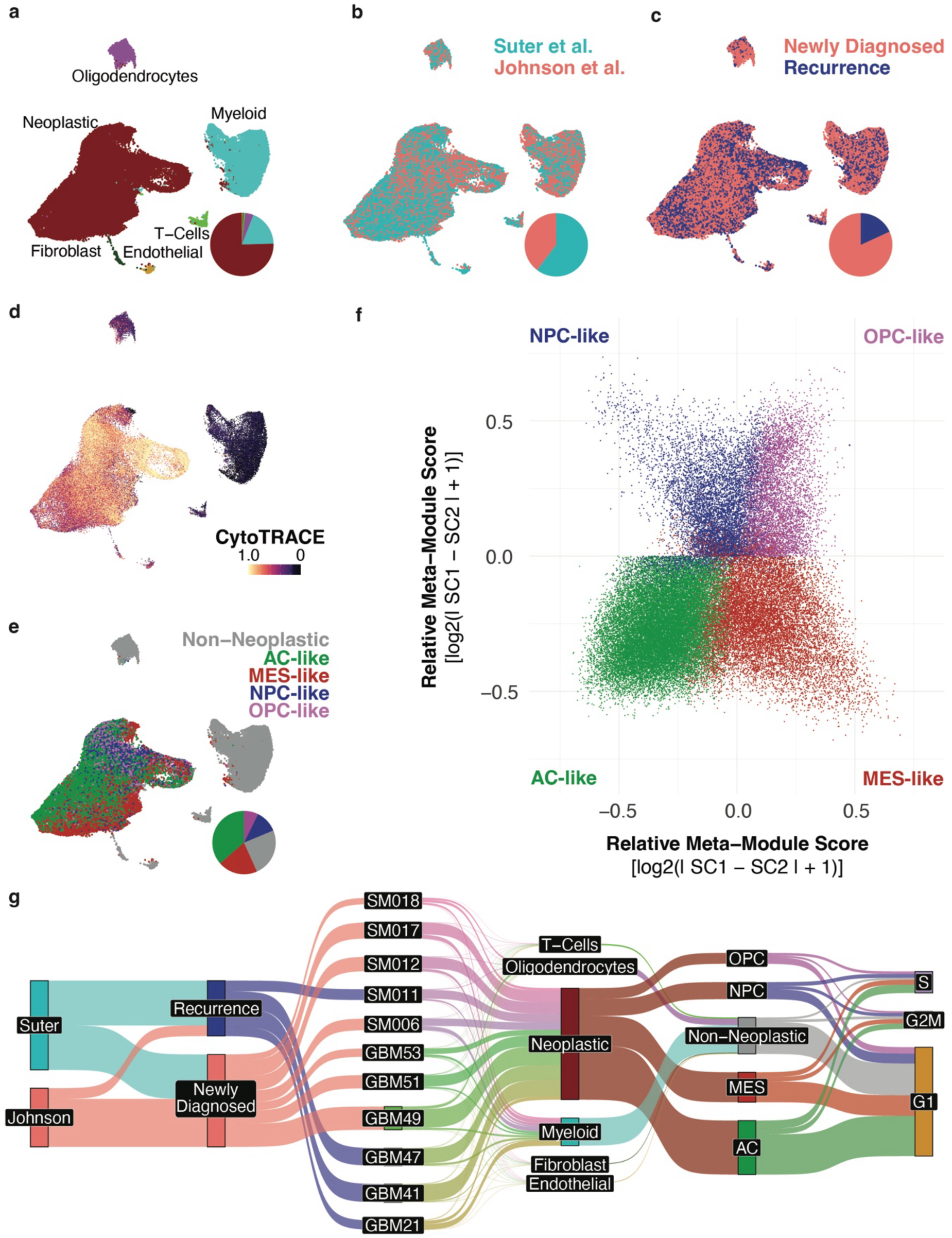
Single-cell RNA sequencing reveals distinct transcriptional states present in newly diagnosed and recurrent GBM. Single-cell RNA sequencing data of 6 patient glioblastoma tumors was integrated and harmonized with an external dataset obtained from Johnson et al. (2021) **a-e.** UMAPs of single-cell transcriptomes colored by **(a)** cell type, **(b)** source dataset, **(c)** whether the tumor was newly diagnosed or recurrent, **(d)** CytoTRACE score, and **(e)** assigned Neftel et al. (2019) GBM cell transcriptional state. **f.** Two-dimensional representation of relative enrichment of GBM cell transcriptional states in neoplastic cells. Cells are colored by assigned transcriptional state identity based on the most predominantly enriched signature of that cell. **g.** Sankey plot depicting proportions of cells grouped by source dataset, occurrence or recurrence, tumor ID, cell type, GBM cell transcriptional state, and expression-based cell cycle phase.

### Integration of L1000 TCSs with expression data of single GBM cells clusters small molecules by mechanism of action

While distinct GBM single-cell transcriptional states have unique disease signatures relative to tumor microenvironment cell controls, this method relies on prior stratification and knowledge of tumor cell populations. We, therefore, sought a way to integrate L1000 TCSs with the expression of individual cells. We utilized the Spearman’s ρ between a single cell’s normalized and scaled expression and TCS signatures to score compounds for their connectivity with each cell in the dataset. Through correlation analysis of each compound’s calculated connectivity across the single-cell landscape, we found that many compounds cluster by their mechanism of action based on their tumor-cell connectivities **(Fig. 2b-c, Supplementary Fig. S2a-b)**. Compound classes such as MEK, PI3K and mTOR, HSP, and HDAC and BET inhibitors cluster individually based on their correlations with GBM tumor single-cell expression data (Pearson’s ρ > 0.7), suggesting that biologically relevant information is retained in this perspective **(Fig. 2c)**. Interestingly, when looking only at 64 FDA-approved oncology drugs identified using annotations from the Drug Repurposing Hub^18^, distinct clusters of small molecules could be identified based on their calculated connectivity with all cells within the GBM scRNAseq atlas. **(Supplementary Fig. S2a-b)**

**Figure 2:**
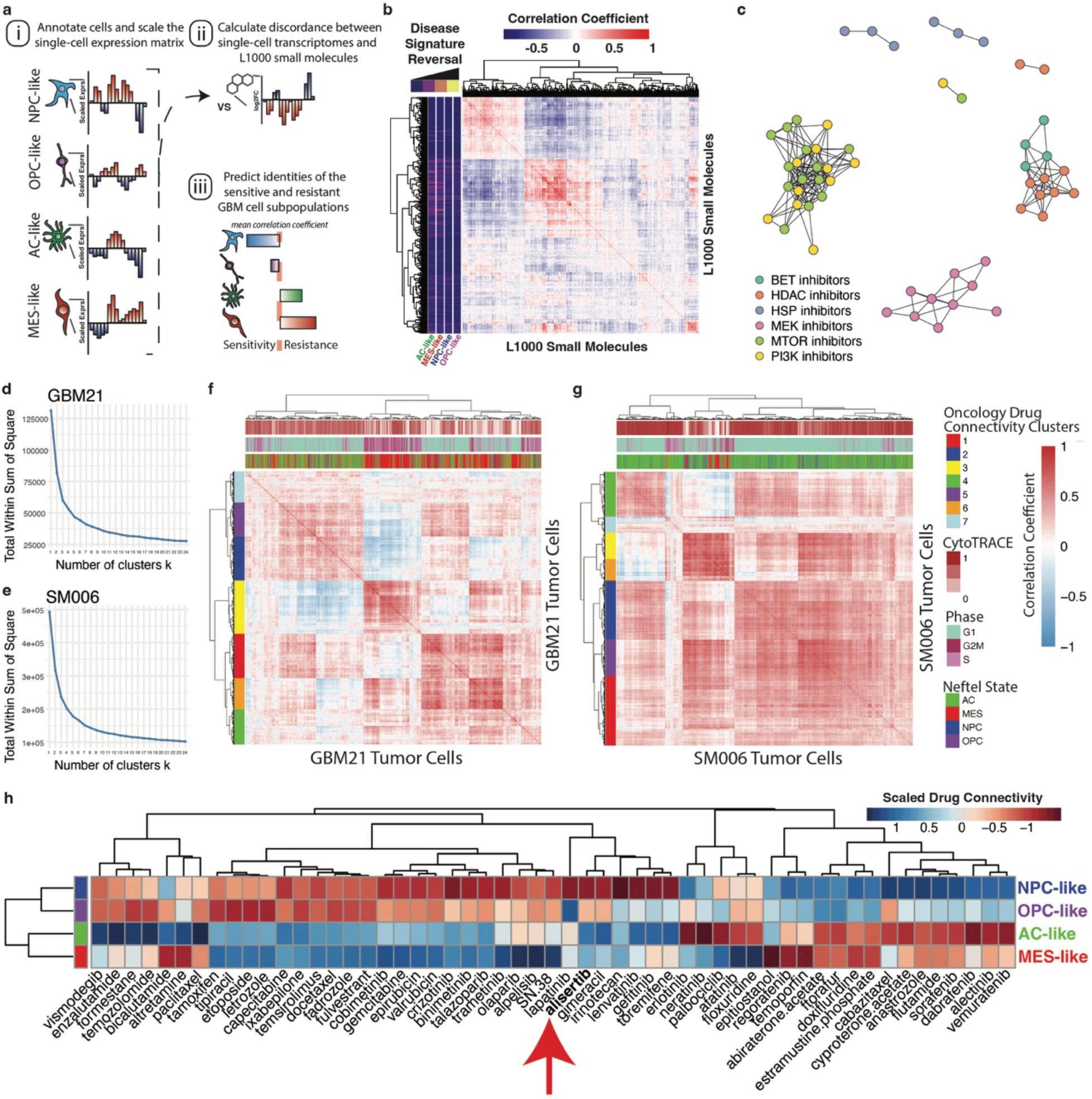
Integration of single-cell expression and small molecule L1000 TCS signatures permits clustering of both compounds and cells by reversal of GBM cell transcriptional state-specific disease signatures. **a.** Schematic of single-cell sensitivity and resistance scoring. **b.** Correlation matrix depicting similarities of L1000 small molecule TCSs by their connectivity to all individual cells within our single-cell atlas. Row annotations depict compound TCS scores for the reversal of AC-, MES-, NPC-, and OPC-like disease signatures. **c.** Network plot of select L1000 small molecules colored by mechanism of action. Connections indicate a Pearson’s ρ > 0.7 between small molecules by their calculated discordances against all single-cells in the GBM dataset as represented in **(b)**. **d.** Elbow plot depicting the within-cluster sum of squares by number of clusters k for GBM tumor cells from patient sample GBM21. **e.** Elbow plot depicting the within-cluster sum of squares by number of clusters k for GBM tumor cells from patient sample SM006 (Johnson et al. dataset). **f-g.** Correlation matrices depicting pairwise Spearman correlations of single GBM tumor cells from individual patient tumor samples (GBM21, in-house) **(f)**, SM006 (Johnson et al.) **(g)** by their connectivity values to 63 FDA-approved oncology drug TCSs. Row annotation bars depict hierarchical clustering identities (k = 7). Column annotations depict CytoTRACE scores, cell cycle phase, and assigned Neftel et al. state identity. **h.** Heatmap showing the scaled inverse single-cell drug connectivities for FDA-approved oncology compounds plus alisertib that were significantly different between Neftel et al. cell states (linear model, BH-adjusted p < 0.05). Heatmap color indicates predicted cell state sensitivity to each compound (red = relative sensitivity, blue = relative resistance).

### The integration of L1000 TCSs of FDA-approved oncology drugs with the expression data of single GBM cells identifies distinct drug connectivity states within single tumors

To investigate whether drug connectivity patterns correlated with previously published GBM cell transcriptional states, we performed a clustering analysis of single cells from individual tumors based on their connectivity to 64 different FDA-approved drugs **(Fig 2d-g)**. Within individual tumors, clear patterns of distinct drug connectivity states are revealed, with some individuals containing at least 7 distinct states. These drug connectivity clusters show patterns in their makeup by Neftel states and developmental potential quantified using CytoTRACE **(Fig. 2f-g, column annotations)**.^17^ Within each individual, cycling cells seem to separate into a unique drug connectivity state.

### Integration of L1000 TCSs with single-cell derived disease signatures predicts differential sensitivity to small molecules

We sought to determine whether the identified pseudo-bulk disease signatures for AC-like, MES-like, OPC-like, and NPC-like states within GBM tumors could be targeted with distinct small molecules or drugs. To accomplish this, we utilized a previously reported method of disease signature reversal scoring.^13^ In brief, a disease-signature-specific discordance score was calculated for all compounds within the L1000 library as the number of genes perturbed by the compound TCS in the opposite direction of the disease signature expression, over the number of genes perturbed by compound TCS in the same direction as in the disease signature **(Fig. 2b row-annotation bar)**. Although a molecule may strongly reverse a bulk expression-based disease signature, reversal may be unique to a subpopulation of tumor cells. Many small molecules are predicted to reverse the disease signatures of MES-like, AC-like, NPC-like or OPC-like cells, but rarely are they predicted to strongly reverse the disease signatures of all transcriptional states **(Fig. 2b, annotation bar, Supplementary Data 4)**.

### In silico perturbation analysis using ISOSCELES predicts the intratumor response to alisertib treatment in vivo

Utilizing our calculated cell-drug connectivity data, we find that GBM cell transcriptional states show differing mean connectivities to different clinical oncology drugs **(Fig. 2h)**. Here, we found 54 out of 64 compounds to have significantly different connectivities between cell states by generalized linear model analysis using *limma* (BH-adjusted p-value < 0.05, **Fig. 2h**), including the aurora kinase inhibitor alisertib. Prior studies reveal that alisertib increases survival in an orthotopic xenograft mouse model of GBM. However, tumors will eventually grow back, suggesting an acquired mechanism of resistance.^13,19–21^ Aside from their role in cell cycle progression, the aurora kinases also play an important role in neurodevelopment and have been extensively studied as targets not only in GBM but many other cancers.^15,21–47^ For this reason, we sought to identify whether a specific cell state conferred resistance to aurora kinase inhibition in GBM. Overall, the alisertib TCS consists of 139 genes, 54 that are upregulated relative to vehicle control, and 85 that are downregulated. Of these 139 genes, 49 were differentially expressed among AC-, MES-, NPC-, and OPC-like cells. Utilizing differential expression between cells belonging to each transcriptional state, we found that the alisertib TCS predicted that alisertib treatment will promote the expression of MES-like specific genes such as HMOX1 or SQSTM1, while reducing the expression of NPC-like specific genes such as CDK4 or EZH2 **(Supplementary Fig. S3b)**. Additionally, the alisertib TCS discordance ratio as utilized in SynergySeq is highest against the NPC-like pseudobulk disease signature **(Supplementary Fig S3c)**. Utilizing the calculated discordances of the alisertib TCS with the individual cells within the dataset, we performed an *in silico* drug connectivity analysis, where cells are predicted to be sensitive or resistant based on their expression concordance with the alisertib TCS **(Fig. 3a-b)**. Predicted alisertib sensitive and resistant cell populations span source dataset, new diagnoses and recurrences, individual patient tumors, and cell cycle phases **(Supplementary Fig. S4a-d)**. Labeling cells by their predominant meta-module enrichment, the predicted resistant cell populations across individual patients consistently showed an increase in proportion of cells within MES-like states and depletion of NPC-like cells **(Fig. 3c)**. The proportion shifts between NPC-like and MES-like states showed the strongest differences, suggesting an NPC-like to MES-like predicted shift **(Supplementary Fig. S3d)**. Differential expression testing between predicted alisertib-resistant and sensitive populations revealed a predicted resistance signature that was most strongly enriched in MES-like cells **(Supplementary Fig. S4f)**. Further, expression of the NPC-like marker CDK4 was significantly decreased in the predicted resistant cell population **(Supplementary Fig. S4e)**.

**Figure 3:**
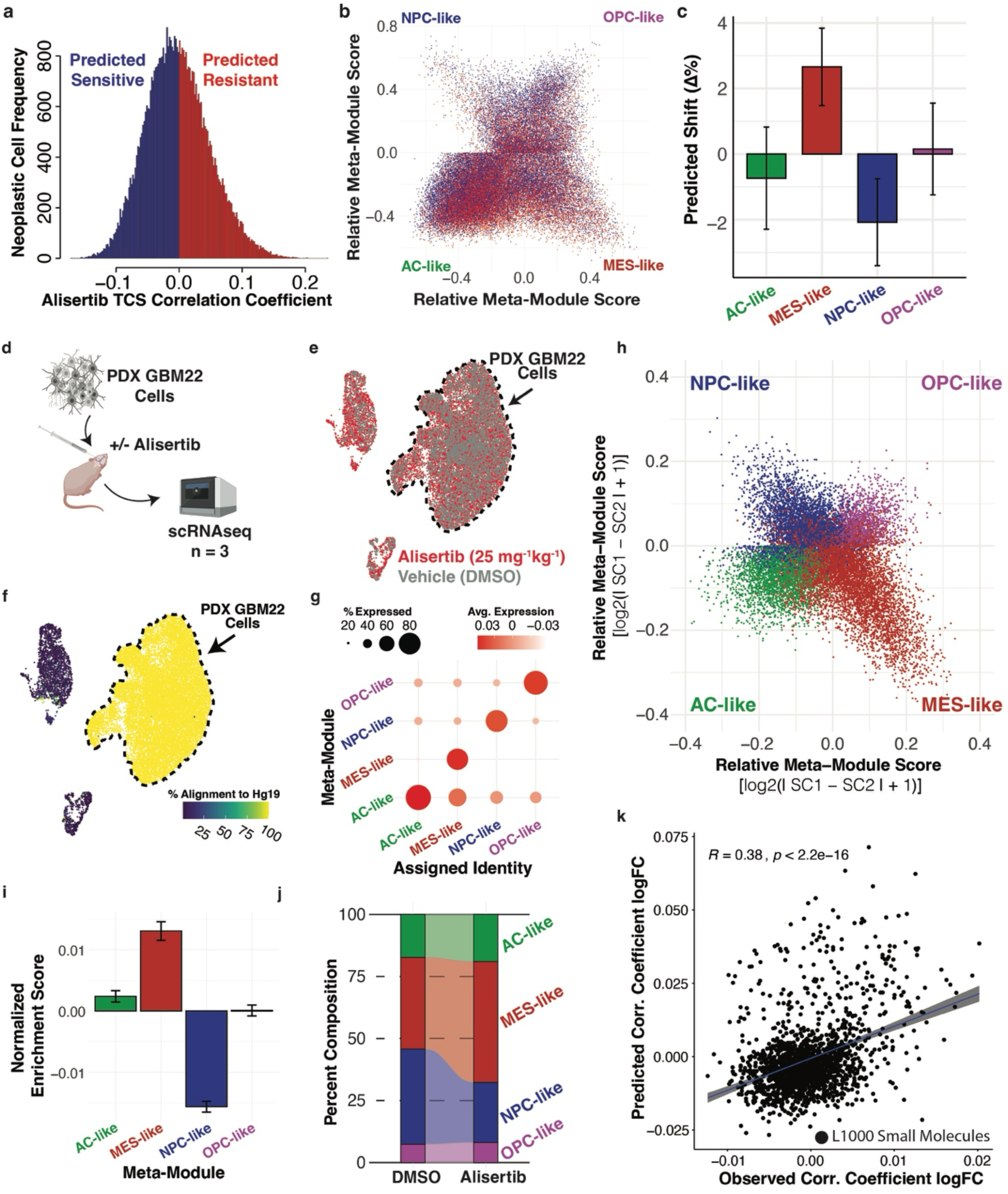
*In silico* perturbation of GBM tumor cell scRNAseq data using an L1000-derived alisertib TCS predicts an NPC-like to MES-like tumor response confirmed *in vivo*. **a.** Histogram of single-cell alisertib TCS correlations (connectivities). Cells with negative correlation coefficients (blue) are predicted to be sensitive to alisertib, while cells with positive correlation coefficients (red) are predicted to be resistant to alisertib. **b.** Hierarchy plot of patient GBM tumor cells colored by predicted sensitivity (ρ < 0) or resistance (ρ > 0) to alisertib. **c.** Bar plot depicting mean shift in proportions of cells in resistant vs. sensitive populations within individual patient tumors. Error bars depict the standard error of the mean percentage differences of individual patient tumors. **d.** Schematic of *in vivo* experiments. **e.** UMAP plot of pre-filter single-cell transcriptomes colored by treatment with either alisertib or DMSO vehicle control. **f.** UMAP of captured cells from orthotopic xenografts colored by percent alignment to the human transcriptome (hg19). **g.** Dot plot of pass-filter, human xenograft single-cell transcriptome expression of Neftel et al. signatures, grouped by assigned transcriptional state identity as assigned based on predominant transcriptional state module expression. **h.** Two-dimensional hierarchical representation of GBM22 xenograft cells’ relative enrichment scores for GBM cell transcriptional state modules. Cells are colored by assigned transcriptional state identity. **i.** Bar plot of mean enrichment shift of transcriptional state signatures in alisertib-treated xenograft cells normalized to DMSO vehicle-treated xenograft cells. Error bars represent 95% confidence interval (ANOVA with Games-Howell Post-hoc, AC-MES p.adj = 3.27e-08, AC-NPC p.adj < 1e-13, AC-OPC p.adj = 3.00e-13, MES-NPC p.adj = 2.88e-08, MES-OPC p.adj = 3.04e-08, NPC-OPC p.adj < 1e-13). **j.** Alluvial plot depicting shift in relative proportion of transcriptional state identities in alisertib and DMSO vehicle control treated xenografts. **h.** Bar plot of change in proportion of cell transcriptional identities in alisertib-treated xenografts relative to DMSO vehicle-treated xenografts. **k.** Scatterplot of L1000 small molecule TCS’s depicting predicted correlation shift (log_2_FC) vs. observed correlation shift (log_2_FC) in alisertib-treated xenografts. Differential small molecule correlations were calculated using *limma*.

To test the predictions from our *in silico* drug connectivity analysis performed using the patient tumor dataset, we utilized scRNAseq of orthotopic xenografts treated with alisertib or vehicle (n=3 per group) to assess alisertib-induced shifts in transcriptional state composition **(Fig. 3d)**. As most tumor cells in our patient dataset were AC-like **(Fig. 1e-g)**, we utilized GBM22 PDX cells, which are transcriptionally classical, for our *in vivo* studies. Single cell analysis of GBM22 tumors isolated from mice revealed all predicted GBM cell states (AC-, MES-, NPC-, and OPC-like) **(Fig. 3g-h, Supplementary Fig. S5a-c)**. Importantly, *in vivo* alisertib treatment induced a shift in expression from NPC-enriched to MES-like transcriptional signatures **(Fig. 3i-j)**, which aligns well with our predictions **(Fig. 3c, Supplementary Figure S3d)** (ANOVA with Games-Howell Post-hoc, AC-MES p.adj = 3.27e-08, AC-NPC p.adj < 1e-13, AC-OPC p.adj = 3.00e-13, MES-NPC p.adj = 2.88e-08, MES-OPC p.adj = 3.04e-08, NPC-OPC p.adj < 1e-13). Likewise, the proportion of GBM22 cells in a predominantly NPC-like state were reduced, while the proportion of GBM22 cells in a predominantly MES-like state were enriched, in line with predictions of proportion shift made within individual patient tumors **(Fig. 3i-j)**.

### *In silico* drug connectivity analysis predicts the intratumor compound discordance shifts observed *in vivo*

To assess whether resistant cell states predicted through *in silico* drug connectivity analysis of the integrated dataset represented those observed to persist through alisertib treatment *in vivo*, we directly compared the differences in global L1000 drug discordance between the predicted alisertib sensitive and resistant cells in the patient data to those observed through comparing alisertib and DMSO treated xenografts **(Fig. 3k, Supplementary Fig. S6a-h)**. We found that predictions of discordance shift from *in silico* drug connectivity analysis with alisertib in patient cells is predictive for certain classes of small molecules, including nucleoside reverse transcriptase inhibitors (Spearman’s ρ = 0.89, p = 0.033), tubulin polymerization inhibitors (Spearman’s ρ = 0.82, p = 0.0068), topoisomerase inhibitors (Spearman’s ρ = 0.82, p = 0.00011), serotonin receptor agonists (Spearman’s ρ = 0.82, p = 0.0068), MEK inhibitors (Spearman’s ρ = 0.77, p = 0.021), HCV inhibitors (Spearman’s ρ = 0.77, p = 0.021), HMGCR inhibitors (Spearman’s ρ = 0.83, p = 0.058), and mTOR and PI3K inhibitors (Spearman’s ρ = 0.94, p = 0.017), **(Supplementary Fig. S6a-h**)).

### Connectivity analysis with an L1000-derived panobinostat TCS separates vehicle-treated from panobinostat-treated cells in both neoplastic and myeloid cell population

To assess the application of our prediction model in *ex vivo* settings, we utilized a publicly available scRNAseq dataset of GBM acute slice cultures treated with the pan-HDAC inhibitor panobinostat (Zhao et al., 2019) **(Supplementary Figure S7a-b)**.^48^ A panobinostat TCS was derived from the 2020 L1000 data release **(Supplementary Figure S7c)**, and used to score panobinostat connectivity of DMSO or panobinostat-treated neoplastic and myeloid cells. In both populations, panobinostat-treated cell populations showed significantly higher panobinostat connectivity values **(Supplementary Figure S7d-e)**. These findings are in agreement with those reported within the original manuscript in which this dataset was reported, wherein panobinostat elicits alterations to the transcriptional states of both tumor and myeloid cell populations^48^.

### Connectivity analysis predicts drug combinations that synergize in vitro and in vivo

Having demonstrated the predictive power of our connectivity analysis framework to identify tumor cells resistant or sensitive to different perturbations, we sought to leverage our framework to identify novel synergistic combinations for glioblastoma **(Fig. 4a)**. First, we utilized a recently published synergy screen performed across 24 independent glioma stem cell lines in spheroid culture to assess the ability of ISOSCELES to predict relative synergy of small molecule combinations **(Supplemental Fig. S8)**.^49^ Using erlotinib, lapatinib, pazopanib and sunitinib as reference compounds **(Supplemental Fig. S8a)**, we ranked 13 small molecule combinations with these inhibitors using an ISOSCELES based combination score **(Supplemental Fig S8b, top heatmap annotation)**. This score is representative of the specificity of a partner molecule in targeting the cell population resistant to the reference molecule and thus reflects the breadth of coverage a combination would have against the GBM cell transcriptional states globally represented within our patient GBM scRNAseq atlas. We demonstrate that this ISOSCELES-based combination score strongly correlates with the mean observed BLISS synergy across cell lines of the combinations present within this screen (Spearman ρ = 0.7, p = 0.01), indicative of the potential that a representative GBM single-cell atlas can be used as a digital twin for identifying more effective combinations **(Supplemental Fig. S8c)**. Next, we focused on the OLIG2 inhibitor CT-179 due to its presumed ability to target OPC-like cells.^50–54^ Using bulk RNA sequencing of GBM8 cells treated with 200nM CT-179 or vehicle, we generated a dose-dependent CT-179 transcriptional signature comprised of 841 differentially expressed genes for analysis of cell-CT-179 connectivity in our patient tumor atlas **(Fig. 4b, Supplementary Fig S9a-c)**. The CT-179 response signature showed enrichment by gene ontology analysis of processes including microtubule and tubulin binding, motor activity, and a strong enrichment for processes related to mitotic cell division **(Supplementary Fig. S9b)**. Using the Spearman correlation of the directional CT-179 response signature with each tumor cell’s scaled expression, cells were binned into predictive sensitive or resistant populations, which spanned all patients within our dataset **(Fig. 4c)**. As expected, the CT-179 predicted sensitive population is primarily OPC-like **(Fig. 4d)**, evident by low CT-179 connectivity in the OPC-like population and high CT-179 connectivity in AC-like cells **(Fig. 4e)**. Using our established workflow for predicting the differential connectivities of L1000 small molecules between predicted sensitive and resistant cell populations, we identified 721 L1000 small molecules that exhibited a significantly reduced connectivity in CT-179 resistant cells compared to CT-179 sensitive cells **(Fig. 5a)**. As above, we also identified 706 discordant L1000 small molecules by their mean connectivity to the CT-179 resistant population **(Fig. 5b)**. 411 L1000 small molecules exhibited both decreased connectivity and overall discordance with the CT-179 resistant population. By weighting the connectivity differentials of these molecules with their mean discordance to the CT-179 resistant cell population (mean Resistant Cell Connectivity, RCC), we established a “combination index score” for each. Top hits by this metric included the EGFR inhibitor varlitinib, as well as docetaxel and indibulin, which elicit anti-neoplastic effects through alteration of tubulin and microtubule assembly and dynamics **(Fig. 5c).**^55–58^ Indeed, EGFR and OLIG2 are expressed by distinct GBM cells in AC-like and OPC- and NPC-like states respectively **(Supplementary Fig. S9d)**. These prioritizations led us towards the use of the anti-EGFR monomethyl auristatin F (MMAF) antibody-drug conjugate Depatux-M (ABT-414), which would target EGFR-expressing cells with MMAF, a tubulin poison, in combination with CT-179. Mice bearing GBM6 orthotopic xenograft tumors were treated with vehicle, CT-179 alone, Depatux-M alone, or CT-179 and Depatux-M in combination. Treatment of mice bearing GBM6 patient derived xenografts with the CT-179/Depatux-M combination increased survival over vehicle treatment or either monotherapy alone **(Fig. 5e)**. In corroboration of the increased survival, bioluminescence imaging reveals a significant reduction in tumor size following treatment with the combination of CT-179 and Depatux-M relative to CT-179 monotherapy alone **(Fig. 5d)**. Collectively, these studies suggest that targeting OPC-like and AC-like cells simultaneously may be advantageous *in vivo*, and that maximizing drug discordance across the tumor cell landscape identifies more effective combinations.

**Figure 4:**
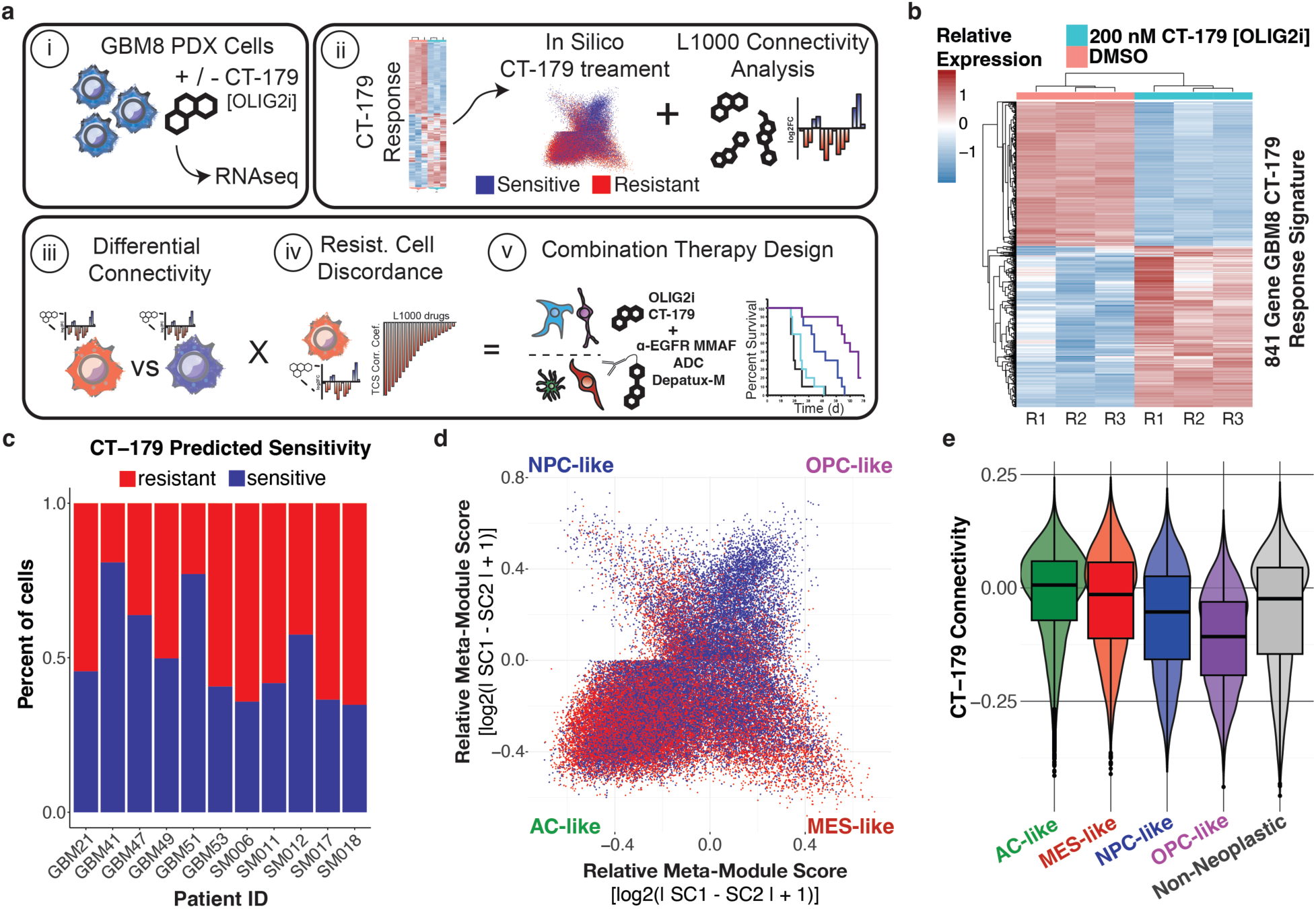
The integration of a bulk-derived transcriptional response signature for the OLIG2 inhibitor CT-179 predicts targeting of OPC-like GBM cells. **a.** Diagram of workflow for identifying CT-179 combinations using ISOSCELES framework. **b.** Heatmap of GBM8 cells treated with vehicle or 200nM CT-179 for 24 hours. Columns represent biological replicates. **c.** Bar plot depicting the proportion of tumor cells within each patient tumor predicted to be sensitive (CT-179 response ρ < 0) or resistant (CT-179 response ρ > 0) to CT-179 treatment. **d.** Hierarchy plot of GBM tumor cells arranged by their relative expression of Neftel et al. states colored by predicted sensitivity or resistance to CT-179 treatment. **e.** Violin and box plot of GBM cell CT-179 connectivity (ρ) grouped by assigned GBM cell transcriptional state.

**Figure 5:**
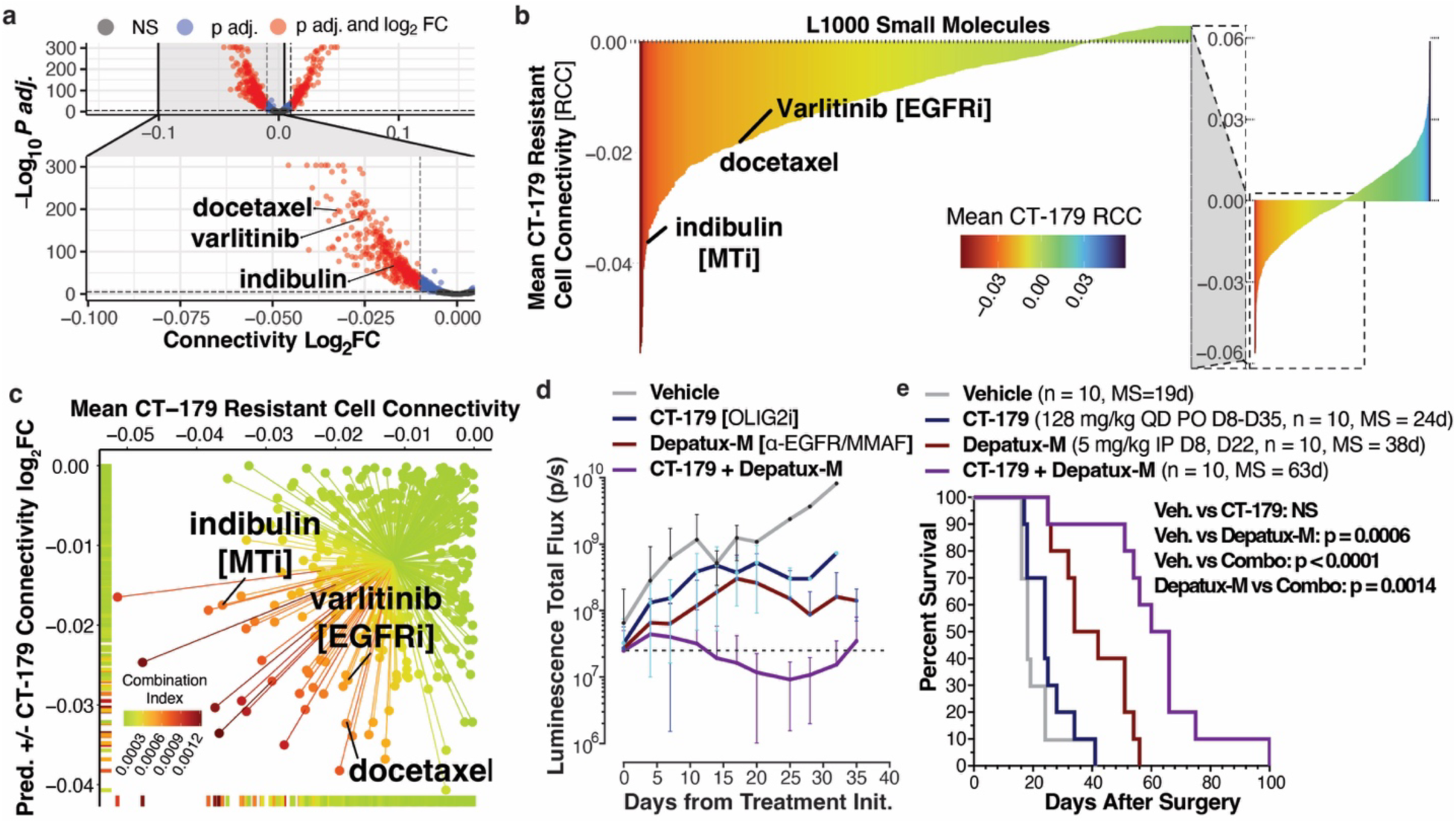
An ISOSCELES combination index predicts synergistic combination of an OLIG2 inhibitor CT-179 with Depatux-M (ABT-414), an anti-EGFR antibody MMAF drug conjugate. **a.** Volcano plot depicting the results from limma-based differential drug connectivity between predicted CT-179 sensitive and resistant GBM cells. Patient ID was used as a covariate in the model design. **b.** Barplot of L1000 small molecule mean resistant cell connectivity (RCC) to predicted CT-179 resistant cells. Color depicts this same value. **c.** Scatterplot of L1000 small molecules plotted by resistant vs. sensitive differential connectivity and mean CT-179 resistant cell connectivity. Colors depict the calculated ISOSCELES combination index, the product of multiplying the differential connectivity log_2_FC values by mean CT-179 resistant cell connectivity values for each individual molecule. Compounds highlighted in (**a), (b), and (c)** are varlitinib, an EGFR inhibitor, and indibulin and docetaxel which act through inhibition of tubulin. **d.** Bioluminescence signal quantification of GBM6-eGFP-FLUC2 orthotopic xenograft tumors (means ± SD, Combo vs Vehicle: p.adj < 0.005 from Day 14, Combo vs. CT-179 monotherapy: p.adj < 0.05 from Day 11, Adjusted p-values from multiple t-tests with Holm–Šídák correction to control the family-wise error rate) **e.** Kaplan-Meier survival curves of mice bearing GBM6-eGFP-FLUC2 orthotopic xenografts treated with indicated therapies (n = 10 per group). MS: median survival. P-values determined using Log-rank (Mantel-Cox) test.

### Release of the ISOSCELES framework as an R package and a web application

We packaged the methodology defined and utilized here into an analytical framework termed ISOSCELES (Inferred cell <u>S</u>ensitivity <u>O</u>perating on the integration of <u>S</u>ingle-<u>C</u>ell <u>E</u>xpression and <u>L</u>1000 <u>E</u>xpression <u>S</u>ignatures), which we have made available as a Shiny web application and as an R package. As we have demonstrated, ISOSCELES integrates bulk drug-response TCS derived from the LINCS L1000 dataset with multi-subject scRNAseq data and facilitates the analysis of drug and cell discordance from multiple perspectives. First, ISOSCELES provides a simple approach to single-cell-derived disease signature generation and permits the analysis of disease signature heterogeneity across tumors relative to non-transformed cell types. Next, ISOSCELES scores small molecules for their reversal potential of the generated disease signatures. Compounds predicted to reverse single-cell-derived disease signatures can then be utilized in ISOSCELES’ *in silico* drug connectivity approach. Here, as demonstrated with alisertib and panobinostat, L1000 TCSs are scored for discordance against the scaled expression of single-cells in a dataset. A custom transcriptional response signature can also be used, as we show in the case of CT-179. Users can then select a sensitivity cut-off (i.e., pseudo-dose), and the single-cell model will be split into predicted sensitive and resistant populations. Characteristics and visualizations, including proportions of annotated identities and predicted drug sensitivities, can then be explored. Additionally, users can identify predicted synergistic compounds through calculation of an ISOSCELES combination index score for each L1000 compound with their drug of interest. All data generated by users of ISOSCELES can be downloaded for ease of access. Single-cell connectivity scores for all compounds against the patient GBM scRNAseq atlas presented in this study are downloadable using the ISOSCELES Shiny web application, which can be found at https://robert-k-suter.shinyapps.io/isosceles alongside instructions for R package download at https://github.com/AyadLab/ISOSCELES **(Fig. 6)**.

**Figure 6:**
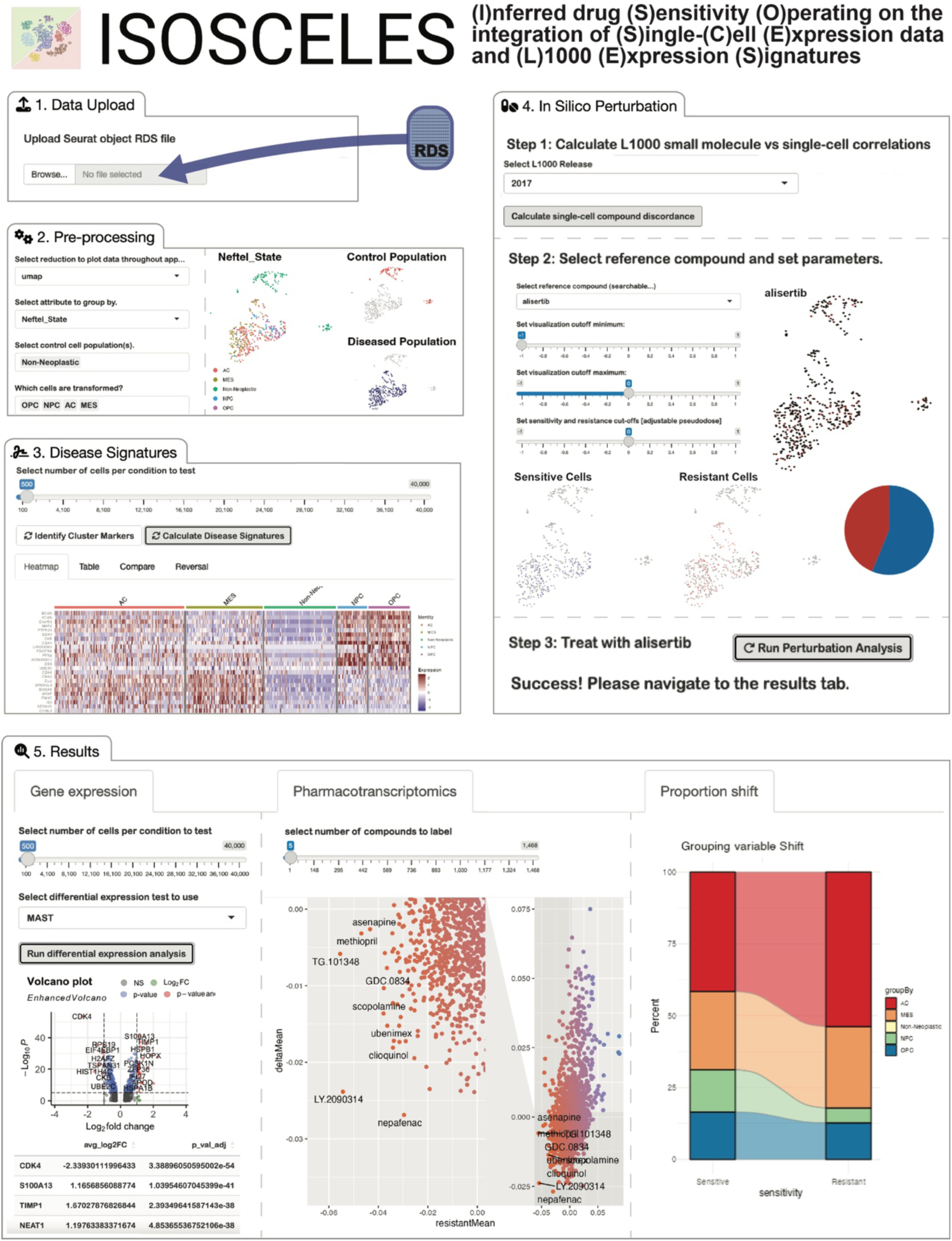
The ISOSCELES framework is available for use as a shiny web application or an R package.

## Discussion

The concepts of disease signature reversal and single-cell omics technologies are becoming a mainstay of drug repurposing and personalized medicine.^11,59–61^ Using principles of disease signature reversal, we have shown that drug response signatures from the LINCS L1000 dataset can be used to identify compounds and compound combinations effective at eliminating specific GBM transcriptional subtypes.^12,13^ Based on these findings, we proposed that if the transcriptional response signatures represent the expression of cells remaining beyond therapeutic pressure, then single cells with expression programs most concordant to that signature would show the highest probability of persistence after treatment and would therefore drive therapeutic resistance.

Here, we describe a novel analytical framework for the identification of small-molecule perturbagens that target distinct transcriptional niches of GBM tumor cells. We created a GBM scRNAseq atlas through the integration of a publicly-available dataset with our novel scRNAseq dataset of newly diagnosed and recurrent patient GBM tumors **(Fig. 1, Supplemental Fig. S1)**. Integration of pharmacological transcriptional response signatures from the LINCS L1000 dataset into the atlas facilitated pharmacotranscriptomic analysis and the development of an *in silico* drug connectivity analysis platform, ISOSCELES, to predict the sensitive and resistant cell populations that would result from specific small molecule treatments **(Fig. 3)**. Importantly, we find that GBM tumor cells can be classified distinctly through their connectivities to current FDA-approved and other clinically relevant oncology drugs **(Fig. 2d-g)**.

Investigating aurora kinase inhibition as a use-case, we used ISOSCELES to predict the sensitivity of NPC-like cells, and the resistance of MES-like cells to treatment with alisertib **(Fig. 3, Supplemental Fig. S3)**. Utilizing single-cell RNA sequencing of orthotopic xenografts treated with alisertib or vehicle, we validated our predictions and demonstrated that MES-like expression and the proportion of predominantly MES-like cells increases with alisertib treatment, while NPC-like expression and the proportion of predominantly NPC-like cells decreased **(Fig. 3, Supplemental Fig. S4)**. ISOSCELES, therefore, has the potential to predict tumor pharmacotranscriptomic shifts occurring under therapeutic pressure **(Fig. 3k)**.

In another use-case, we utilized a publicly available single-cell dataset of GBM acute slice cultures treated with the HDAC inhibitor panobinostat to reinforce that L1000-derived TCS connectivity is predictive of treatment response at single-cell resolution **(Supplemental Fig. S7)**. Importantly, we found that cell-panobinostat TCS connectivity predicts single-cell resolution alterations in both the tumor and myeloid cell populations of the tumor microenvironment (TME), as reported in the original publication **(Supplemental Fig. S7)**.^48^ These findings should be explored deeper in the future to assess the potential of ISOSCELES-based prioritization of small molecules targeting distinct TME cell phenotypes.

Acquired treatment resistance is common in glioblastoma, which is notoriously heterogeneous and plastic at single-cell resolution.^6–8,10,62–64^ Longitudinal studies in glioblastoma implicate epigenetic intratumor heterogeneity in standard-of-care treatment resistance.^65–67^ Transcriptional subtype switching occurs often upon GBM recurrence.^10,67^ Thus, the characterization of the transcriptional response of acquired therapy resistance and the identification of means by which to circumnavigate this resistance is essential. Importantly, others have shown that the most discordant cells within a tumor are predictive of patient outcome in the clinic and, further, that predictors of treatment response based on single-cell RNA sequencing data outperform those based on bulk expression profiles.^59^ They also demonstrate that the predictive power of the data lies within the most ‘resistant’ cells in the dataset. Here, we posit that this means that identifying drugs that target these ‘resistant’ cells in the context of a reference treatment can identify more effective therapies.

We evaluate this hypothesis by using our combination scoring approach on our patient scRNAseq atlas and a bulk RNA-seq derived transcriptional response signature for the OLIG2 inhibitor, CT-179. CT-179 is a new small molecule recently characterized as a potential therapeutic for other brain cancers, such as medulloblastoma.^50–54^ OLIG2 has also been characterized as a key dependency in GBM, regulating a neurodevelopmental transcriptional program key in gliomagenesis and maintenance of different tumor cell states.^68,69^ However, targeting of OLIG2 alone is not sufficient in the face of GBM intratumor heterogeneity. As expected, cell and CT-179 connectivity predict the targeting of OPC-like GBM cells **(Fig. 4e)** and an AC-like CT-179 resistant state. In line with this observation, combination indexing with ISOSCELES highlighted EGFR and tubulin inhibitors (varlitinib, indibulin, docetaxel) to combine with CT-179. In support of these predictions, Depatux-M (ABT-414), an antibody-drug-conjugate consisting of an EGFR antibody conjugated to MMAF, a tubulin inhibitor, synergizes with CT-179 to attenuate tumor growth and to extend survival *in vivo* **(Fig. 5d-e)**.^70^

While other tools exist for exploring and utilizing the L1000 CMap datasets, such as the L1000CDS2 or TargetTranslator, ISOSCELES is a novel tool with distinct functionality and uses.^71,72^ The L1000CDS2 serves as a browser for identifying single datapoints in the data correlated with input gene signatures, but does not facilitate sample-by-sample analysis, integration with standard single-cell workflows, nor an implementable combination prioritization strategy.^71^ TargetTranslator is a powerful framework for identifying druggable targets using gene expression data, but it does not inform on what the resistant cell states to inhibiting these targets would be, or which targets should be engaged simultaneously.^72^ Importantly, ISOSCELES permits analyses utilizing L1000-derived TCSs as well as custom input response signatures, a feature that can be used to characterize novel drugs in the context of similarity to other L1000 molecules, predicted target cell populations, and prioritization of additional combinations.

### Limitations of the study

Although we have developed a novel computational platform to predict drug combinations based on scRNAseq information, we do acknowledge that there are limitations to our work. ISOSCELES, as well as any other predictive platform, is reliant on the data it is built upon. For one, most of the input for ISOSCELES is from LINCS, and therefore, compounds have to have been profiled using the L1000 assay. To temper this limitation, we added a novel feature to allow investigators to add their own transcriptional consensus signatures from RNAseq data of treated cells, which should expand the utility of ISOSCELES. Additionally, ISOSCELES uses scRNAseq data as its input, and both cell types (e.g., neurons) and spatial architecture are lost during the processing of samples. Therefore, the integration of single-cell gene expression with other diverse data types, such as DNA methylation, epigenetic profiling of histone acetylation, proteomics, and spatial transcriptomics, could enhance and refine the predictive power of ISOSCELES into a more comprehensive prediction model. Similarly, it may be possible to use different reference “normal” cells to enhance concordance and discordance predictions for tumor cells in ISOSCELES with the addition of the aforementioned data. Exploring the utility of single-nuclei RNA-seq (snRNAseq) datasets, for example, may provide both additional non-tumor cells and different tumor cell states not captured by scRNAseq. Future studies are required to compare scRNA and snRNAseq predictions from ISOSCELES and evaluate the capacity to integrate the data they provide.

Importantly, the response signatures themselves used to calculate compound discordance with single cells could be optimized further. Many alternate approaches to TCS generation have been validated.^71,73^ Additionally, gene expression altered over longer periods compared to the time points assessed within the L1000 dataset (e.g., 24 hours), may be more representative of long-term alterations that would occur in situ in GBM as well as other diseases. Lastly, the selection of the most tissue-relevant cell lines from among those profiled within the L1000 dataset may improve the predictive power for certain tissues. Although we used all perturbed L1000 lines as opposed to GBM lines only, we have demonstrated in a prior publication the general applicability of L1000 TCSs derived from non-GBM cell lines to predict efficacy in GBM and other brain cancer types.^12,13,74^

## Conclusions

Collectively, our studies demonstrate that the application of disease signature reversal to the transcriptional profiles of single cells can predict the sensitive and resistant cell populations within GBM tumors to aid the selection of therapeutic compound combinations for the treatment of patients with GBM. This highlights the potential utility of single-cell transcriptomics for future clinical trial design, wherein patients could be stratified based on the abundance of specific cell populations within tumors. Further, ISOSCELES is in line with dynamic precision medicine initiatives, where intratumor dynamics in response to therapies can be forecasted to inform effective therapeutic combinations or series of therapeutics that maintain effective targeting of a fluid tumor cell landscape. Lastly, the ISOSCELES platform could be leveraged to investigate the potential to reverse distinct cellular phenotypes to more desirable ones in other disease types. All analyses outlined are easily accessible through the ISOSCELES R package and application. ISOSCELES is released with an easy-to-use graphical user interface developed on the R Shiny platform, facilitating its use on pre-processed datasets to users without coding backgrounds, which increases accessibility to a broader range of pharmacology domain experts and represents a significant advance towards data-driven personalized medicine.

## Methods

### Experimental model and study participant details

#### Collection of newly diagnosed and recurrent GBM tumors

Six (6) patients with GBM permitted tissue collection before tumor resection under approved IRB study 20170887 (MODCR00002196). Of the 6 enrolled patients, 3 were newly diagnosed with GBM and 3 had tumor recurrence after previously receiving standard of care treatment with temozolomide chemotherapy and radiotherapy. The 3 newly diagnosed patients had not received any treatment before resection. After surgical resection, University of Miami Hospital pathologists separated the resected tissue, which was then transferred from the operating suite to the laboratory on ice. Tissue from the enhancing edge was then used for dissociation and single-cell RNA sequencing (see below).

#### GBM cell xenograft models

Human patient-derived GBM cells (GBM6, GBM8, GBM22) were obtained from the Mayo Clinic Brain Tumor Patient-Derived Xenograft National Resource. GBM22 cells were transfected to express GFP as previously described, while GBM6 cells were transfected to express eGFP-FLUC2 as previously described.^70^ GBM8 was derived from a newly diagnosed tumor in a female patient and is molecularly IDHwt, Classical, EGFR amplified, and MGMT methylated. GBM22 was derived from a newly diagnosed tumor in a male patient and is molecularly IDHwt, Classical, MGMT methylated. GBM6 was derived from a newly diagnosed tumor in a male patient and is molecularly IDHwt, with EGFRVIII mutation, MGMT unmethylated. Orthotopic xenografts

#### Mouse models

Nu/Nu mice from Charles River Laboratories were utilized for *in vivo* experiments. Orthotopic implantations took place when mice were 6-8 weeks old. All procedures were approved by the University of Miami IACUC, protocol number 18-014. All animal procedures were performed per guidelines and regulations, including ARRIVE guidelines, and performed as previously described.^13,70,75,76^

### Method details

#### Preparation of newly diagnosed and recurrent GBM tumors for scRNAseq

We performed single-cell RNA sequencing on the 6 patient GBM tissue samples collected at the time of surgical resection (n = 3 newly diagnosed, n = 3 recurrent, **Fig. 1**) After resection, University of Miami hospital pathologists separated the GBM tissue, which was transferred from the operating suite to the laboratory on ice. Tumor samples were dissociated using the Worthington Biochemical Papain Dissociation System.^77^ Tumor tissue was finely minced with a sterile surgical blade and incubated in sterile Earl’s Balanced Salt Solution (EBSS) containing 20 units of papain per mL in 1 mM L-cysteine with 0.5mM EDTA and DNase for 45 minutes at 37 °C while shaking. Following papain incubation, the tissue was gently triturated with a 5 mL serological pipette until it was adequately dissociated. The resulting suspensions were pelleted and resuspended in EBSS with ovomucoid protease inhibitor, bovine serum albumin (BSA), and DNase. This suspension was layered on top of ovomucoid protease inhibitor solution with BSA to form a gradient, which was centrifuged at 70xg for 6 minutes. The supernatant, which contained cellular debris, was removed. The pellets were resuspended, and the gradient centrifugation was repeated. The resulting pellet was then washed 2 times with ice cold PBS with 0.1% BSA, passed through a 0.45 μM cell strainer, and resuspended in ice cold PBS with 0.1% BSA. This resulting suspension was then checked for cell viability by staining an aliquot with acridine orange and propidium iodide (AO/PI) and counted on a Nexcelom K2 Cellometer. All samples had cell viability greater than 90% prior to loading of the single-cell chips. Single cells were captured using a 10X Genomics Chromium Controller, and patient single-cell gene expression libraries were prepared using 10X Genomics 5’ Gene Expression chemistry. All libraries were sequenced on an Illumina NextSeq500.

#### Pre-processing and quality control of patient-derived scRNA-seq data

Raw scRNA-seq dataset was aligned and demultiplexed using CellRanger (10X Genomics), and the resulting filtered unique molecular identifier (UMI) matrices were used for downstream analysis in the R package Seurat (v4).^78–82^ Within Seurat, poor quality cells were further filtered by removing outliers based on the number of detected UMIs and unique transcripts, as well as percent expression of mitochondrial transcripts. Pass-filter cells were those with greater than 200 and less than 4500 detected features, and with less than 12.5% mitochondrial expression. Data from 42,839 pass-filter single-cell transcriptomes across 33,469 features were normalized and variance stabilized using a regularized negative binomial regression implemented within the R package scTransform, and individual patient datasets were integrated using an anchor-based technique utilizing the Pearson residuals.^78–82^

#### Harmonization and integration of in-house and external scRNAseq datasets

Previously published single-cell RNA sequencing data from Johnson et al. (2021) were downloaded from Synapse (https://synapse.org/singlecellglioma). The five IDHwt GBM samples out of the eleven total glioma samples available in the dataset were used.^16^ To integrate the Johnson et al. dataset with our internal dataset, each was first split into individual datasets based on the patient source. Each individual patient’s expression matrix was then independently prepared for integration using scTransform, and integrated together using the anchor-based technique in Seurat **(Fig. 1b)**. 28,130 single-cell transcriptomes from Johnson et al. were integrated with our dataset for a total of 70,969 single cells. Integrated expression values were used for initial shared-nearest-neighbor (SNN) clustering, dimensionality reduction, and cell type identification **(Fig. 1a)**. Downstream analysis including differential expression and enrichment scoring was performed using log-normalized counts.

#### Generation of GBM cell transcriptional state-specific disease signatures

SNN clustering on the patient-derived, variance-stabilized, integrated scRNAseq data identified distinct clusters of neoplastic and non-neoplastic cells. To confirm the identification of these neoplastic cells, we assessed transcriptional diversity with the R package CytoTRACE using raw counts as input. CytoTRACE scores cells based on their developmental potential, reflecting the plasticity of GBM tumor cells relative to the TME **(Supplementary Fig. S1)**.^17^ Non-neoplastic cell identities were assigned by expression of canonical cell type markers identified through differential expression with MAST (Model-based Analysis of Single-cell Transcriptomics) **(Supplementary Fig. S1, Supplementary Data 1)**.^83^ The neoplastic cells were further subdivided and assigned GBM cell state identities using GBM cell transcriptional state signatures previously reported in Neftel et al. (2019) and the R package singscore **(Supplementary Fig. S1)**.^6,84^ Briefly, the raw count matrices across all samples were log-normalized and scaled, regressing out S-Phase and G2/M-Phase expression module scores as calculated in Kowalczyk & Tirosh et al. (2015). These scaled data were then used to generate gene rank lists for each individual cell. Using the R package *Singscore*, gene set enrichment analysis was performed on each cell independently for the six Neftel et al. GBM cell state signatures (meta-modules). The means of NPC1- and NPC2-like, and MES1- and MES2-like state enrichment scores were calculated and represented as NPC-like and MES-like enrichment, respectively. Each cell’s assigned transcriptional identity reflects its predominant, most-enriched GBM cell state expression program (AC, MES, NPC, OPC). Thus, the identities assigned within this manuscript indicate the predominant transcriptional program of that cell. Hierarchy plots were generated using the R package scrabble.^6^ Differential expression testing using MAST was performed between each of the state-specific clusters and all non-neoplastic cells to generate disease signatures **(Fig. 2a, Supplementary Data 2)**.^83^ MAST was used to account for the technical pitfall of stochastic dropout and the resulting bimodal expression distributions inherent in scRNAseq. Individual patient signatures were generated by comparing each individual’s tumor cells to all non-neoplastic cells from all patients within the dataset **(Supplementary Data 3)**. Heatmaps were generated using the R package pheatmap.^85^

#### In silico drug connectivity analysis with the aurora kinase inhibitor, alisertib

The overlap of the alisertib TCS and results from differential expression testing using MAST between patient neoplastic cells grouped by predominant transcriptional state enrichment was calculated to identify potential state-specific expression perturbed by alisertib (Fig. 4a). Single-cell Spearman’ ρ values for the alisertib TCS were extracted from the L1000 small molecules versus single-cell correlation matrix. Neoplastic patient cells were divided into predicted alisertib resistant and alisertib sensitive populations wherein cells concordant (Spearman’s ρ > 0) with the alisertib TCS were identified as a predicted resistant population, and those discordant (Spearman’s ρ < 0) with the alisertib TCS were identified as a predicted sensitive population **(Fig. 3a-b)**. The predicted shift in cell states was then calculated, demonstrating an increase in MES-like cells and a decrease in NPC-like cells **(Fig. 3c)**. Proportions across source dataset, initial occurrence or recurrence, source patient, and cell cycle phase were also compared **(Supplementary Fig. S3a-d)**. Proportion shifts of cells predominantly in each transcriptional state within each individual were compared using ANOVA with a Games-Howell post-hoc test. Resulting adjusted p-values were used to visualize the relative probability of predicted directional state abundance shifts **(Supplementary Fig. S3d)**. Differential expression testing between predicted resistant and sensitive populations was performed using MAST, and cells were scored for their enrichment of overexpressed genes using Seurat’s AddModuleScore function **(Supplementary Fig. S4f)**.

#### GBM cell orthotopic xenograft, alisertib treatment, and scRNAseq library creation

Human patient-derived GBM cells (GBM22) expressing GFP obtained from the Mayo Clinic Brain Tumor Patient-Derived Xenograft National Resource were implanted in the brains of nude mice as previously described **(Fig. 3d)**.^19,70,76^ Cells were visualized to confirm GFP expression prior to intracranial implantation into mice. Briefly, 6-8 weeks old Nu/Nu mice (Charles River Laboratory) were anesthetized with ketamine (100 mg/kg) and xylazine (10 mg/kg), and a hole was drilled through the skull (1 mm anterior and 2 mm lateral to bregma). Using a Hamilton syringe, 150,000 GBM22 cells suspended in PBS were injected stereotactically at a depth of 2 mm. The skin incision was closed using surgical glue with sutures or wound clips and mice were given buprenorphine as analgesic. Procedures were approved by the University of Miami IACUC, protocol number 18-014. All procedures were performed in accordance with guidelines and regulations, including ARRIVE guidelines.

Following tumor implantation, mice recovered for a period of 7 days. GBM22 orthotopic tumor-bearing mice were treated intraperitoneally with either Alisertib (25mg/kg) (drug formulation: 5% DMSO / 30% PEG400 / 5% Tween-80 / 60% PBS) or vehicle control 5 days per week and monitored daily for signs of cognitive decline and body weight (BW). Tumors were collected at the time of initial signs of cognitive decline or 15% of BW loss.

Tumor cells from three individual mice within each treatment group were captured and pooled for single cell RNA-sequencing. Tumor-bearing mice were perfused with PBS, and their brains were removed. Tumor was isolated and dissociated using the Worthington Biochem Papain dissociation kit. Cells were washed two times using ice-cold PBS with 0.1% BSA (w/v), then checked for viability using AO/PI dye and a Nexcelom K2 Cellometer. Cells from the dissociation suspension were loaded into a 10X Genomics’ Chromium chip using 10X Genomics’ NextGEM 3’ chemistry targeting the capture of approximately 10,000 cells per sample. With each individual animal experiment, paired-end library sequencing of DMSO and alisertib-treated single-cell libraries was performed in tandem on a single S1 NovaSeq flow cell, with asymmetric read lengths as recommended by 10X Genomics.

#### Pre-processing and quality control of in vivo scRNAseq data

Sequencing basecalls were input into CellRanger V3.0.2 for demultiplexing and fastq generation. Alignment and gene quantification were first performed in tandem to a dual-species reference transcriptome for both hg19 (human) and mm10 (mouse), obtained from 10X Genomics (refdata-cellranger-hg19-and-mm10-3.0.0.). Using the R package Seurat (V4.0.1), the percentage of raw UMI counts within each cell aligning to either mm10 or hg19 transcriptomes was calculated and used in tandem with SNN clustering to separate out human GBM22 cells from contaminating mouse cells **(Fig. 3e)**. GBM22 cell barcodes were then used to isolate these human cells from the data aligned to the GrCH38 (human) transcriptome for downstream gene expression analysis **(Fig. 3f)**. Outlier cells were removed based on UMI count, detected gene count, and percent of mitochondrial transcript counts. Cells with less than 200 or more than 6000 detected genes, or with more than 10% of mitochondrial genome alignment were removed for downstream analysis. Pass-filter data was comprised of 16,102 GBM22 single-cell transcriptomes with 10,394 measured features.

#### Generation of a CT-179 GBM cell transcriptional response signature

GBM8 cells (Primary, IDHwt, Female, Classical, EGFR amp, MGMT methylated) were obtained from the Mayo Clinic Brain Tumor Patient-Derived Xenograft National Resource. GBM8 cells were treated with 100nM CT-179, 200 nM CT-179, or vehicle control, and RNA was extracted following a 24 hour treatment period. RNA sequencing was performed as previously described.^86^ Differential expression between 200nM CT-179 and vehicle-treated GBM8 cells was performed using DESeq2 **(Supplementary Fig. S9a)**. Differentially expressed genes retained within the signature showed a logFC greater than 0.425, calculated as two standard deviations from the mean of all logFC values, and an adjusted p-value less than 0.05 **(Fig 4b)**. Gene ontology analysis was performed on this filtered signature using clusterProfiler **(Supplementary Fig. S9b)**.^87–91^ Enrichment of the signature in 100nM CT-179 treated, 200nM CT-179 treated, or vehicle-treated GBM8 samples was calculated using the R package Singscore and compared to confirm the dose-dependence of our signature **(Supplementary Fig.S9c)**.

#### In vivo evaluation of CT-179 and Depatux-M in combination

Orthotopic implantation of GBM6-eGFP-FLUC2 was performed as previously published.^70^ One week following surgery, mice were treated with vehicle, CT-179, Depatux-M, or a combination of CT-179 and Depatux-M (n = 10 per group). CT-179 was formulated in 0.5% methylcellulose, 0.5% Tween80 in water, and stored at 4C. Fresh batches were made on the first day of dosing, and on day 18 of dosing. CT-179 formulation was stirred at RT for 30 minutes prior to use, and was delivered at a dose of 128mg/kg QD PO from days 8-35. Depatux-M (ABT-414) was formulated in PBS, and was delivered at a dose of 5 mg/kg IP on day 8 and day 22. Tumor formation was confirmed, and tumor volume was assessed using bioluminescence imaging (BLI) **(Fig. 5d).** BLI was performed as previously reported using an IVIS 2.50 cooled charged-coupled device camera (Xenogen 200 series) and Living Image software.^70^ Mice were dosed with 10 mg/kg Cycluc1 by intraperitoneal injection and imaged 10 minutes later under isoflurane anesthesia.^70^ A moribund state was defined by outward neurological deficits (altered gait, hunched posture, circling, poor body condition score). Survival differences were assessed using the Log-Rank test and Kaplan-Meier analysis **(Fig. 5e)**.

### Quantification and statistical analysis

#### Ranking of L1000 compounds against GBM cell transcriptional state disease signatures

The L1000 assay contains 978 “landmark” genes that can be measured to algorithmically infer the expression of 11,300 additional genes.^71,73,92^ Using a previously published algorithm to rank discordance of L1000 perturbations, small molecules’ response signature discordance with scRNAseq-derived disease signatures were calculated for each compound in the dataset.^13,14,73,92^ This disease-specific discordance score was calculated for all compounds within the L1000 library as a ratio of the number of genes with discordant expression (opposite gene expression direction/sign) between the transcriptional consensus signatures (TCS) and the cell state disease signature over the number of genes with concordant expression (same gene expression direction/sign) between the TCS and the disease signature **(Annotation Bar Fig. 2b, Supplementary Data 4)**. ^13^

#### Ranking of single compounds against individual cell expression programs

The filtered single cell counts matrix output by CellRanger was log normalized and scaled using the R package Seurat. Using the normalized consensus signatures from the LINCS L1000 2017 dataset, predicted relative sensitivity of each single-cell to each L1000 perturbation (connectivity) was measured as the Spearman’s ρ between a compound’s consensus signature and each individual cell’s scaled gene expression of the L1000 genes present in each compound’s consensus signature **(Fig. 2a)**. Pearson correlation analysis was applied to the compound versus cell coefficient matrix to visualize small molecule clustering based on integration with the single-cell data **(Fig. 2b)**. Mechanism of action annotations were obtained from the Broad Institute Drug Repurposing Hub **(Supplementary Fig. S2)**.^18^ Single-cell drug connectivities were compared between cell states using a generalized linear model implemented through limma. The mean connectivities of significantly different FDA approved oncology compounds in addition to alisertib (p < 0.05) for each cell state were then scaled and visualized by heatmap **(Fig. 2h)**.

#### Tumor cell drug connectivity clustering analysis

Using the Drug Repurposing Hub, 63 FDA-approved oncology drugs with L1000-derived TCSs from the 2017 data release were identified. A pair-wise Spearman correlation matrix was generated on the calculated drug connectivity values of all tumor cells from each patient in the single-cell atlas for these drugs. Euclidean distance was calculated using the base R function dist().Ward’s hierarchical clustering of the distance matrix was performed using the base R function hclust(). Within cluster variance (sum of squares) was visualized for each number of resulting clusters K using the R package factoextra.^93^ K of 7 was selected for representative visualizations based on the resulting elbow plot.

#### Xenograft tumor cell stratification and state shift analysis

Xenograft tumor cells were assigned GBM cell state identities using GBM cell transcriptional state signatures (Neftel et al., 2019), as described above for patient data **(Fig. 3g-h, Supplementary Fig. S4b-c)**. Singscore derived enrichment scores and proportions of cells predominantly within each transcriptional state were compared between DMSO vehicle and alisertib-treated xenografts **(Fig. 3i-j)**. The analysis pipeline is available at https://github.com/AyadLab/ISOSCELES-dataProcessing.

#### Patient and xenograft pharmacotranscriptomic comparisons

To investigate the relationship between cells in GBM22 xenografts and those captured directly from patient tumors, GBM22 cells from DMSO treated xenografts were input into ISOSCELES, and were scored for correlation with all small molecule TCSs. The mean compound-cell connectivity values (Spearman’s ρ) were calculated across each transcriptional state identity. Using Spearman’s ρ once more, these connectivity values were compared to the same mean connectivity values obtained using patient neoplastic cells **(Supplementary Fig. S6a)**. To assess whether ISOSCELES predictions of alisertib resistant and sensitive cell identities from patient data adequately modeled the pharmacotranscriptomic changes occurring *in vivo,* we utilized a linear model to calculate global L1000 small molecule discordance shift across both patient and xenograft datasets. A pseudo-count of 1 was added to all values in the compound-cell connectivity matrices, which were then fit to a linear model using *limma* and *voom.* Patient-based compound-cell connectivity values were grouped by predicted resistance or sensitivity, and differential discordance was calculated across populations within each individual patient tumor. The xenograft cell discordance differential was calculated in the same manner. Resulting logFC values for each compound were compared across *in silico* and *in vivo* analyses using Pearson’s ρ **(Supplementary Fig. S5b) (Supplementary Data 5)**.

#### Panobinostat TCS connectivity analysis in ex vivo GBM acute slice culture

Single-cell RNA sequencing data of 5 paired acute slice culture samples derived from patient glioblastoma resections treated with either DMSO vehicle or the HDAC inhibitor Panobinostat (0.2 μM) was obtained from GEO accession GSE148842.^48^ Single-cell transcriptomes with less than 200 detected genes or with greater than the 75^th^ percentile of reads detected, calculated respectively per sample, were kept for downstream analysis. The pass-filter dataset was comprised of 62,250 single-cell transcriptomes with 58,828 features. Individual samples were then integrated using reciprocal PCA within Seurat. SNN clustering and differential expression analysis were used to identify discrete cell types within the integrated dataset. A TCS was generated for panobinostat utilizing L1000 data from within the 2020 release as described above **(Supplemental Fig. 6c).** Cell connectivity to the panobinostat TCS was calculated across all cells within the dataset. A Wilcoxon rank sum test with continuity correction was utilized to compare panobinostat connectivity across treatment groups within the determined neoplastic (37,682 cells) and myeloid cell (13,017 cells) populations respectively, and was visualized by violin and area-normalized kernel density estimate plots created with ggplot2 **(Supplemental Fig. 6d-e)**.

#### Combination scoring analysis

We determined the potential to apply ISOSCELES connectivity analysis in ranking synergistic small molecule combinations. An ISOSCELES combination score was implemented as the differential logFC of each small molecule’s connectivity between cells predicted to be sensitive or resistant to a reference small molecule, multiplied by that small molecule’s mean connectivity to cell population predicted to be resistant to the reference small molecule. To assess the utility of this approach, synergy screen data was obtained from Houweling et al, 2023 supplementary data.^49^ Raw sum synergy BLISS scores from short-term treatment assays of 24 different cell lines in spheroid culture were used. Following filtering of the synergy screen data for compounds present within the L1000, 4 potential reference small molecules with more than two associated combinations with molecules also in the L1000 data were used for analysis. These reference molecules were erlotinib, lapatinib, pazopanib, and sunitinib. Neoplastic cells from the patient single-cell atlas were scored for connectivity of these and all partner molecules within the screen. Connectivity cut-offs for sensitive or resistant binning of the patient neoplastic cells was calculated as the mean of all cells’ connectivity for each reference molecule respectively. Using limma, the differential connectivity across reference molecule sensitive and resistant cell populations, for each of the partner compounds present in the screen, were calculated for each individual patient within the atlas. The resulting logFC values from each patient for each partner molecule were averaged across all individual patients. These aggregate differential connectivities were then multiplied by the mean connectivity score of each respective small molecule to the resistant cell population to the respective reference small molecule across all patients. These scores were then scaled between 0 and 1. Calculated combination score and observed BLISS raw sum synergy for each combination was compared using Spearman correlation in each of the 24 cell lines tested, and across the aggregate mean BLISS scores for all cell lines for each combination.

#### Scoring of L1000 small molecules for predicted synergy with CT-179

CT-179 single-cell connectivity was calculated as the Spearman correlation between the normalized and scaled expression cells within our dataset against the log_2_FC calculated for 200nM CT-179 treatment of GBM8 cells. As above, sensitive and resistant cells were defined as those having CT179 connectivities less than and greater than 0 respectively. L1000 drug-cell connectivities were transformed using the Fisher’s Z transformation, and limma was used to calculate the differential connectivities to L1000 molecules between predicted CT-179 sensitive and resistant cells, using patient ID as a latent variable. Compounds with logFC < 0.5 and FDR < 0.05 were retained. The logFC values for these compounds were then weighted by multiplying by their respective mean resistant cell-drug connectivities to derive our ISOSCELES-based combination index. Thus, a more positive combination index value indicates both decreased connectivity to that drug in the CT-179 reference population and an overall discordance of the drug with CT-179 predicted resistant cells. This combination index can be defined as the following:

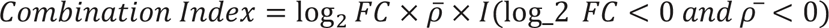

Where a combination index is only calculated when both the differential connectivity (log_2_FC) and the mean resistant cell connectivity (RCC) (ρ) are negative values. This combination index is a feature implemented within the ISOSCELES R package and associated shiny GUI.

## Additional resources

ISOSCELES Shiny Web GUI: https://robert-k-suter.shinyapps.io/isosceles ISOSCELES R package: https://github.com/AyadLab/ISOSCELES

## Data availability

All data needed to generate the conclusions in this manuscript are present in the paper and/or the Supplementary Materials. Our generated single-cell data of patient GBM tumors, including processed Seurat objects, are available via the Gene Expression Omnibus (GEO) with accession ID GSE229779. Xenograft single-cell data including processing Seurat objects are also available via the Gene Expression Omnibus (GEO) with accession ID GSE231489. These GEO series are under GEO SuperSeries GSE231490 (Reviewer Token ujyvuussjrkrfcr). Additional patient single-cell data from Johnson et al. was obtained from https://synapse.org/singlecellglioma, and acute slice culture single-cell data was obtained from GEO at GSE148842.

## Code availability

Data, code, and materials are available to any researcher, and processing pipelines can be found at https://github.com/AyadLab/ISOSCELES-dataProcessing or within the ISOSCELES package source code available at https://github.com/AyadLab/ISOSCELES, which contains a comprehensive readme file, tutorials, and example datasets for use with the R package-based application.

## ACKNOWLEDGEMENTS

The authors thankfully acknowledge Dr. Corneliu Sologon, Dr. Jenny Kemper, Marissa Brooks and Yoslayma Cardentey of the Sylvester Comprehensive Cancer Center Oncogenomics Shared Resource (OGSR). We thank the Georgetown University Lombardi Cancer Center, the Georgetown University Department of Veterinary Resources, the Georgetown University Information Services, the University of Miami Department of Veterinary Resources, and the Miami Project to Cure Paralysis. We thank the Mayo Clinic Brain Tumor Patient-Derived Xenograft National Resource. We thank Luke Tallon and Lisa Sadjewicz of the University of Maryland Institute for Genome Sciences. We thank Dr. Vance Lemmon, Dr. John Bixby, Dr. Claes Wahlestedt, Dr. William J. Harbour, Dr. Daniel Pelaez, Dr. David J. Robbins, and all members of the Schürer and Ayad laboratories for invaluable discussions of these studies.

## Author contributions

*Conceptualization:* R.K.S., A.J., V.S., and N.G.A.; *Methodology, investigation, visualization:* R.K.S.; Tissue Acquisition: M.E.I, R.J.K.; *scRNAseq:* R.K.S., S.L.W; In vitro studies: A.W.; Computational design: R.K.S.; *Data analysis:* R.K.S., R.V., S.K., M.D., M.S.,L.Z.; P.P. *App development:* R.K.S., A.J., R.V., M.D., G.B., L.R.; *Animal studies:* R.K.S., A.J., S.H.H., S.K., W.W., M.C., D.B., E.B.R., A.O., M.G.A.; *Writing and revision:* R.K.S., A.J., S.K., R.V., M.D., M.I.D, M.E.I., R.J.K., G.S., S.K.; N.G.A.; *Supervision:* S.C.S., N.G.A:

## Supplementary Materials

**Supplementary Data 1:** Comma-separated table of MAST differential expression testing results between discrete cell types identified in patient GBM scRNAseq data.

**Supplementary Data 2:** Comma-separated table of MAST differential expression testing results between GBM tumor cells within each Neftel et al. transcriptional state and non-tumor cell types captured from the tumor microenvironment.

**Supplementary Data 3:** Comma-separated table of MAST differential expression testing results between GBM tumor cells within each Neftel et al. transcriptional state, within each individual patient tumor, respectively, compared to all non-tumor cell types captured from the tumor microenvironment.

**Supplementary Data 4:** Comma-separated table of calculated reversal scores for each compound in the L1000 dataset against each of the aggregate (across all patients) disease signatures calculated for each Neftel et al. GBM cell transcriptional state.

**Supplementary Data 5:** Excel .xlsx file of results of differential drug connectivity between alisertib-resistant and sensitive cell populations determined *in silico* (Tab 1: patient_DrugDiscordancelimma_result) and *in vivo* (Tab 2: pdx_DrugDiscordancelimma_result).

