## Supplemental Figures for "Drug and Single-Cell Gene Expression Integration Identifies Sensitive and Resistant Glioblastoma Cell Populations"

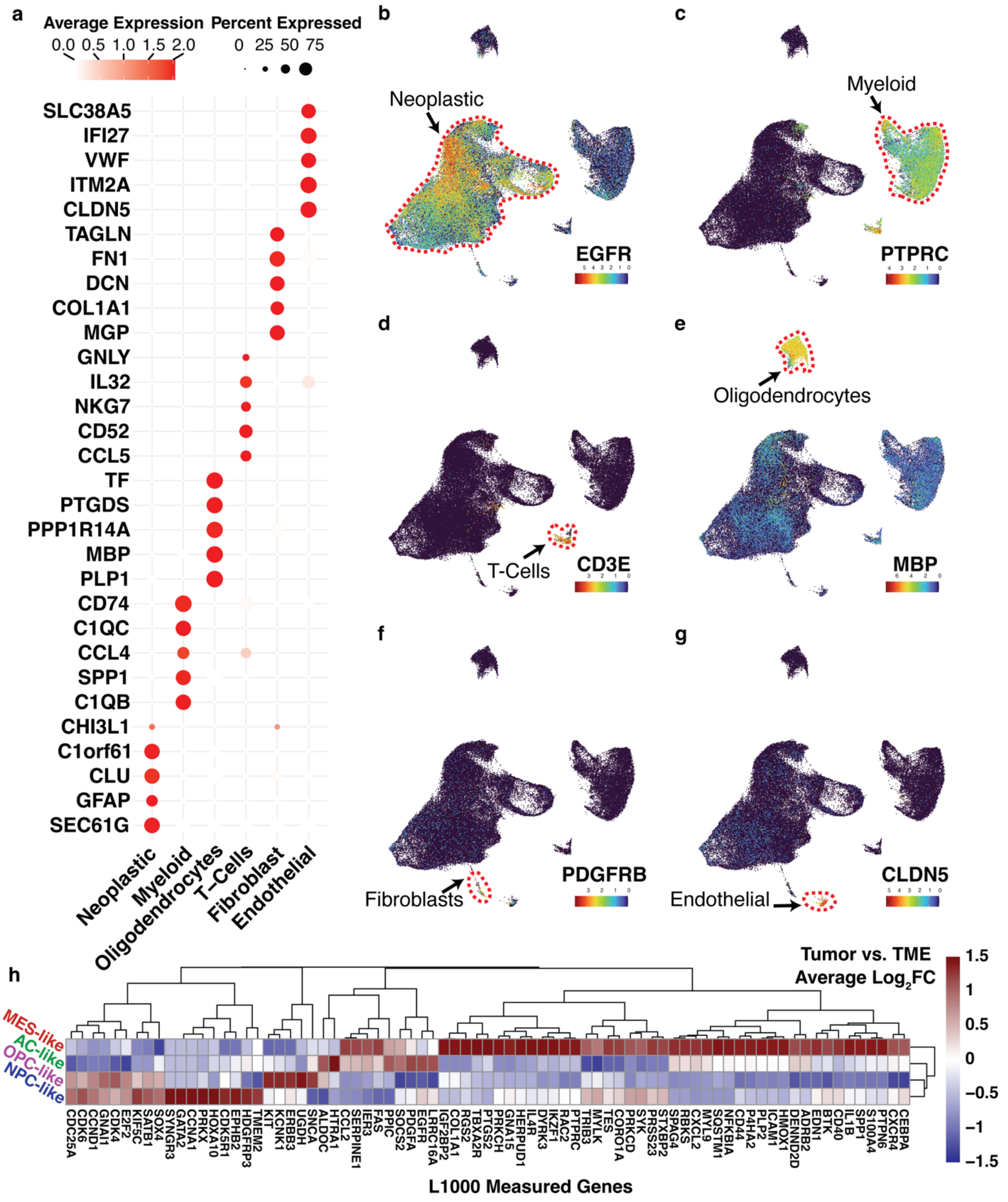
a
Average Expression
0.0 0.5 1.0 1.5 2.0
Percent Expressed
0 25 50 75
SLC38A5
IFI27
VWF
ITM2A
CLDN5
TAGLN
FN1
DCN
COL1A1
MGP
GNLY
IL32
NKG7
CD52
CCL5
TF
PTGDS
PPP1R14A
MBP
PLP1
CD74
C1QC
CCL4
SPP1
C1QB
CHI3L1
C1orf61
CLU
GFAP
SEC61G
Neoplastic
Myeloid
Oligodendrocytes
T_Cells
Fibroblast
Endothelial
b
Neoplastic
EGFR
c
Myeloid
PTPRC
d
T-Cells
CD3E
e
Oligodendrocytes
MBP
f
Fibroblasts
PDGFRB
g
Endothelial
CLDN5
h
Tumor vs. TME
Average Log2FC
1.5
1
0.5
0
−0.5
−1
−1.5
MES-like
AC-like
OPC-like
NPC-like
CDC25A CDK6 CCND1 GNAI1 CDK4 E2F2 KIF5C SATB1 SOX4 SYNGR3 GATA2 CCNA1 PRKX HOXA10 CDK5R1 EPHB2 HDGFRP3 TMEM2 KIT KCNK1 ERBB3 UGDH SNCA ALDOC HTRA1 CCL2 SERPINE1 IER3 FAS PPIC SOCS2 PDGFA EGFR LRRC16A IGF2BP2 COL1A1 RGS2 TBXA2R PTGS2 PRKCH GNA15 HERPUD1 IL4R DYRK3 IKZF1 RAC2 PTPRC TRIB3 MYLK TES CORO1A PRKCD SYK PRSS23 STXBP2 SPAG4 RBKS CXCL2 MYL9 SQSTM1 NFKBIA CD44 P4HA2 PLP2 ICAM1 HMOX1 DENND2D ADRB2 EDN1 BTK CD40 IL1B SPP1 S100A4 PTPN6 CXCR4 CEBPA
L1000 Measured Genes

**Supplementary Figure S1: Single-cell RNA sequencing reveals distinct non-neoplastic cell types present in both newly diagnosed and recurrent GBM. a.** Dot plot of top marker expression of discrete cell types. The dot size encodes the percentage of cells within each class (cell type) with detected expression, while the color depicts the mean expression level across all cells within each class (cell type). **b-g.** UMAP plots depicting the scaled expression of the cell type-specific markers EGFR **(b)**, PTPRC **(c)**, CD3E **(d)**, MBP **(e)**, PDGFRB **(f)**, and CLDN5 **(g)**. **h.** Heatmap depicting relative differential expression of L1000-measured genes within pseudo-bulk disease signatures calculated using MAST. Heatmap color represents the scaled $log_2FC$ relative to each GBM transcriptional state, calculated for each transcriptional state vs non-neoplastic cell populations within the scRNAseq data.

a

b

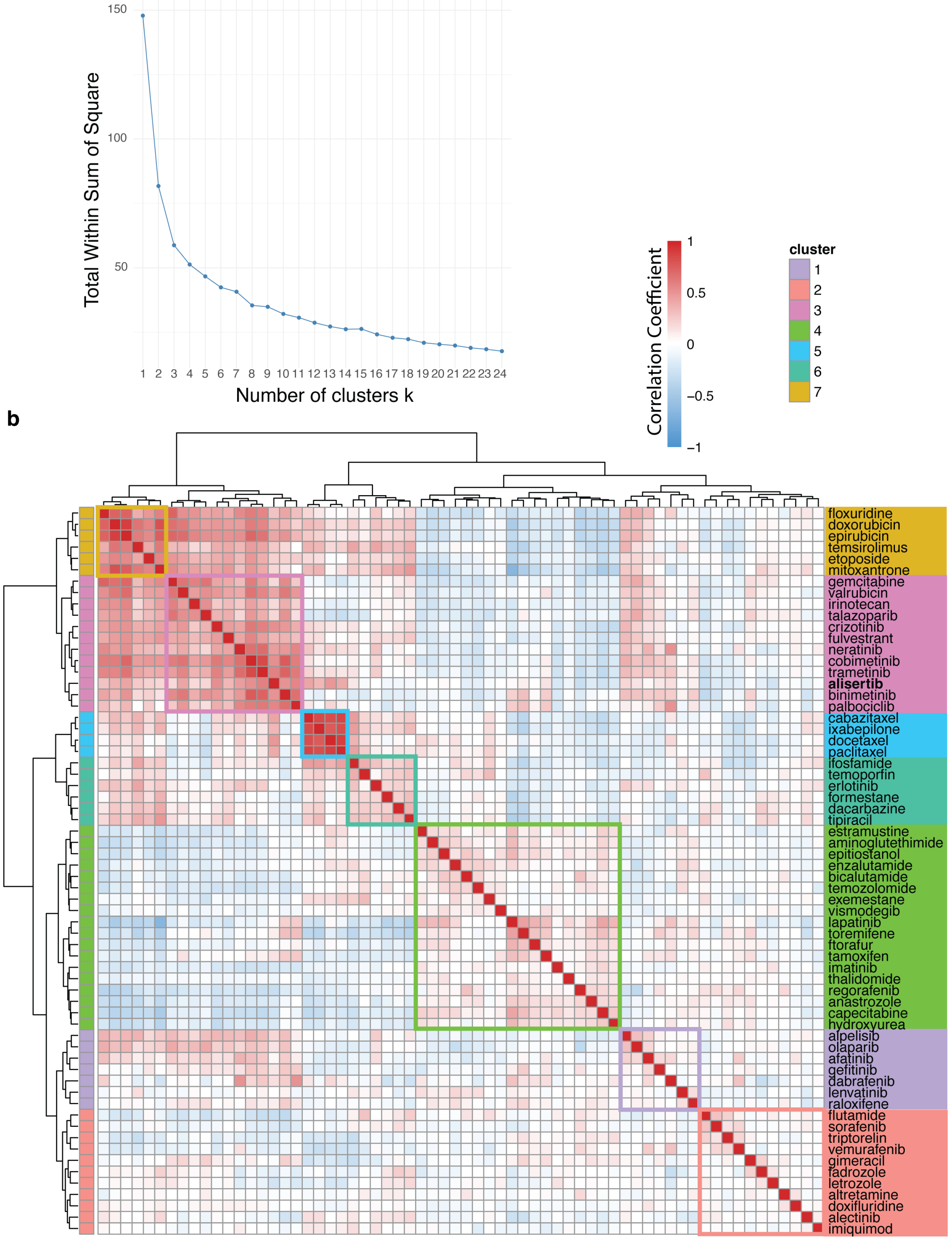

Total Within Sum of Square
Number of clusters k
Correlation Coefficient
cluster
floxuridine
doxorubicin
epirubicin
temsirolimus
etoposide
mitoxantrone
gemcitabine
valrubicin
irinotecan
talazoparib
crizotinib
fulvestrant
neratinib
cobimetinib
trametinib
alisertib
binimetinib
palbociclib
cabazitaxel
ixabepilone
docetaxel
paclitaxel
ifosfamide
temoporfin
erlotinib
formestane
dacarbazine
tipiracil
estramustine
aminoglutethimide
epitiostanol
enzalutamide
bicalutamide
temozolomide
exemestane
vismodegib
lapatinib
toremifene
ftorafur
tamoxifen
imatinib
thalidomide
regorafenib
anastrozole
capecitabine
hydroxyurea
alpelisib
olaparib
afatinib
gefitinib
dabrafenib
lenvatinib
raloxifene
flutamide
sorafenib
triptorelin
vemurafenib
gimeracil
fadrozole
letrozole
altretamine
doxifluridine
alectinib
imiquimod

**Supplementary Figure S2: FDA-approved oncology drugs can be clustered on their L1000 TCS connectivity to GBM tumor cell expression. a.** Elbow plot depicting the within-cluster sum of squares by number of clusters k for connectivity values of 64 clinical oncology drug TCSs with all GBM tumor cells from 11 patients in the scRNAseq atlas. **b.** Correlation matrix heatmap depicting pairwise Spearman correlations of 64 clinical oncology drugs based on their connectivity values to all GBM tumor cells from the 11 patients in the scRNAseq atlas.

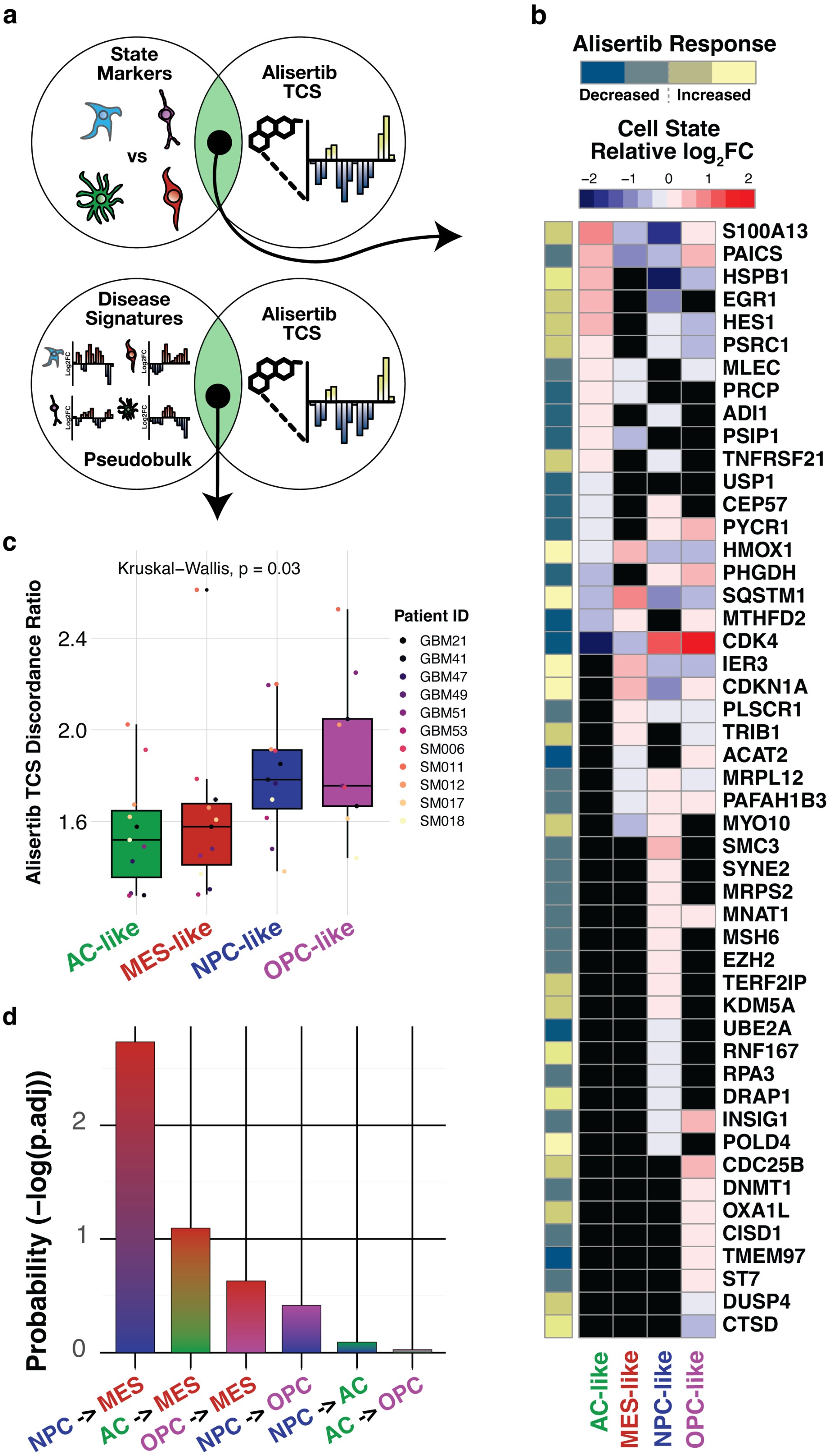

a
State Markers
vs
Alisertib TCS
Disease Signatures
Pseudobulk
b
Alisertib Response
Decreased
Increased
Cell State Relative log2FC
−2 −1 0 1 2
S100A13
PAICS
HSPB1
EGR1
HES1
PSRC1
MLEC
PRCP
ADI1
PSIP1
TNFRSF21
USP1
CEP57
PYCR1
HMOX1
PHGDH
SQSTM1
MTHFD2
CDK4
IER3
CDKN1A
PLSCR1
TRIB1
ACAT2
MRPL12
PAFAH1B3
MYO10
SMC3
SYNE2
MRPS2
MNAT1
MSH6
EZH2
TERF2IP
KDM5A
UBE2A
RNF167
RPA3
DRAP1
INSIG1
POLD4
CDC25B
DNMT1
OXA1L
CISD1
TMEM97
ST7
DUSP4
CTSD
AC-like
MES-like
NPC-like
OPC-like
c
Kruskal−Wallis, p = 0.03
Alisertib TCS Discordance Ratio
2.4
2.0
1.6
Patient ID
GBM21
GBM41
GBM47
GBM49
GBM51
GBM53
SM006
SM011
SM012
SM017
SM018
AC-like
MES-like
NPC-like
OPC-like
d
Probability (−log(p.adj))
2
1
0
NPC -> MES
AC -> MES
OPC -> MES
NPC -> OPC
NPC -> AC
AC -> OPC

**Supplementary Figure S3: Interrogation of an alisertib TCS using single-cell RNA sequencing indicates the predicted sensitivity of an NPC-like GBM cell state. a.** Diagram describing interrogation of the L1000-derived alisertib TCS using GBM cell state markers as in (**b**) or by discordance ratio with GBM cell state disease signatures as in (**c**). **b.** Heatmap depicting differential expression of cell state markers that overlap with alisertib TCS genes. Cell color indicates the relative $log_2FC$ of each gene, and black heatmap cells indicate a gene was not differentially expressed by a given cell state. Row annotation depicts magnitude and direction of gene expression changes of the alisertib TCS. **c.** Box plot of cell state disease signature discordance to the alisertib TCS by individual patient tumor. Cell state disease signatures were calculated using MAST to compare cells within each GBM cell state to non-neoplastic cell types within each individual. **d.** Bar plot depicting the pairwise post-hoc -log(p.adj) values of proportion change for each state combination as calculated from (**c**).

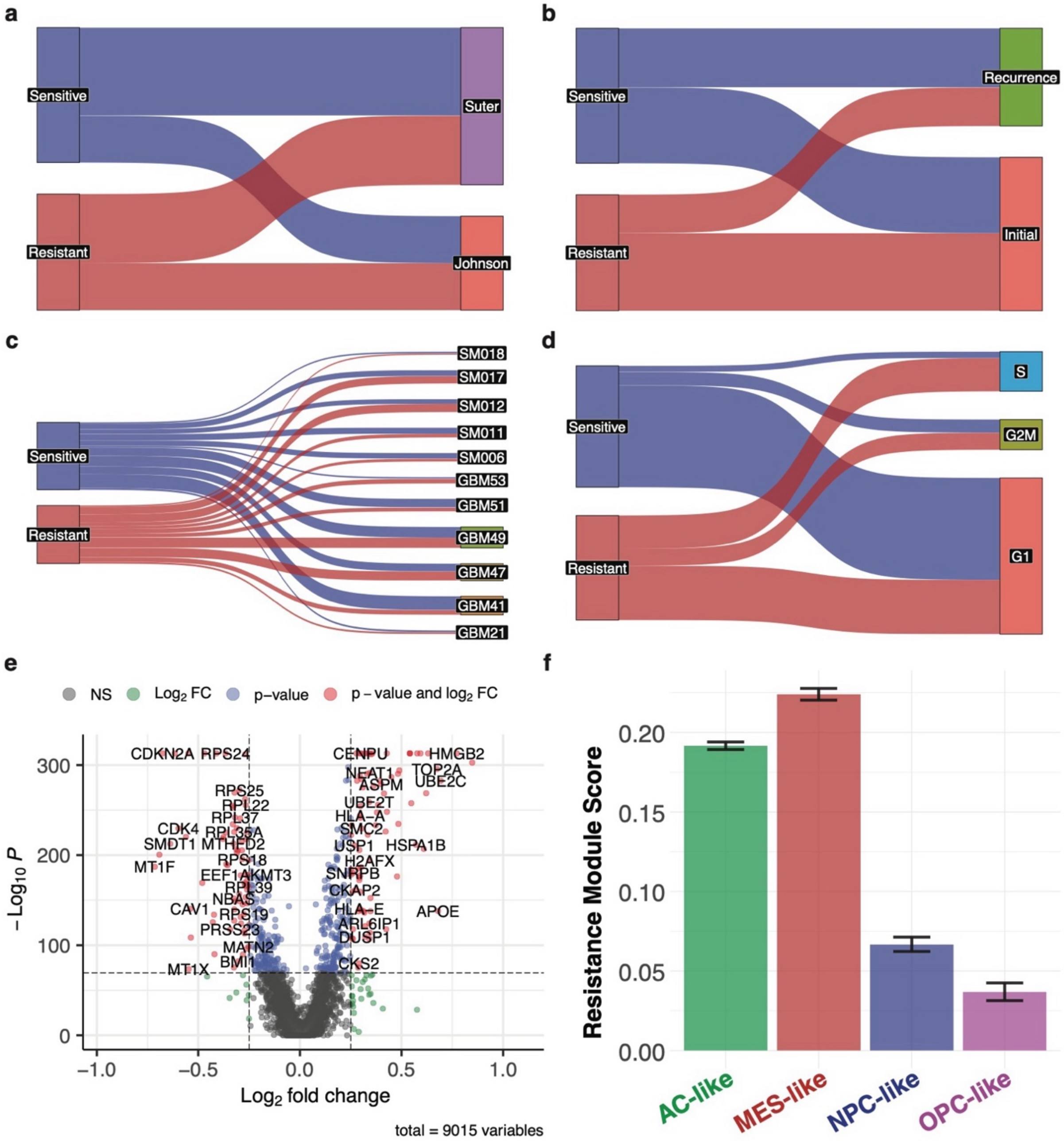

a
Sensitive
Resistant
Suter
Johnson
b
Sensitive
Resistant
Recurrence
Initial
c
Sensitive
Resistant
SM018
SM017
SM012
SM011
SM006
GBM53
GBM51
GBM49
GBM47
GBM41
GBM21
d
Sensitive
Resistant
S
G2M
G1
e
NS
Log2 FC
p−value
p − value and log2 FC
− Log10 P
Log2 fold change
total = 9015 variables
CDKN2A RPS24
CENPU
HMGB2
TOP2A
UBE2C
NEAT1
ASPM
UBE2T
HLA−A
SMC2
USP1
HSPA1B
H2AFX
SNRPB
CKAP2
HLA−E
ARL6IP1
DUSP1
CKS2
APOE
RPS25
RPL22
RPL37
CDK4
RPL35A
SMDT1
MTHFD2
RPS18
MT1F
EEF1AKMT3
RPL39
NBAS
CAV1
RPS19
PRSS23
MATN2
BMI1
MT1X
f
Resistance Module Score
AC-like
MES-like
NPC-like
OPC-like

**Supplementary Figure S4: ISOSCELES predicts an alisertib-resistant cell population that spans the source dataset, initial occurrence or recurrence, individual patients, and cell cycle phase. a-d.** Sankey diagrams depicting proportions of predicted sensitive and resistant cell populations across **(a)** source dataset, **(b)** initial occurrence or recurrence, **(c)** individual patients, and **(d)** cell cycle phase. **e.** Volcano plot of differentially expressed genes between ISOSCELES-predicted resistant and sensitive cell populations. Genes with positive $\log_2$FC represent those overexpressed in ISOSCELES-predicted resistant tumor cells. **f.** Bar plot of mean module scores for ISOSCELES-predicted resistant cell overexpressed genes as depicted in **e** of patient neoplastic cells grouped by predominant GBM cell transcriptional state. Error bars represent the 95% confidence interval.

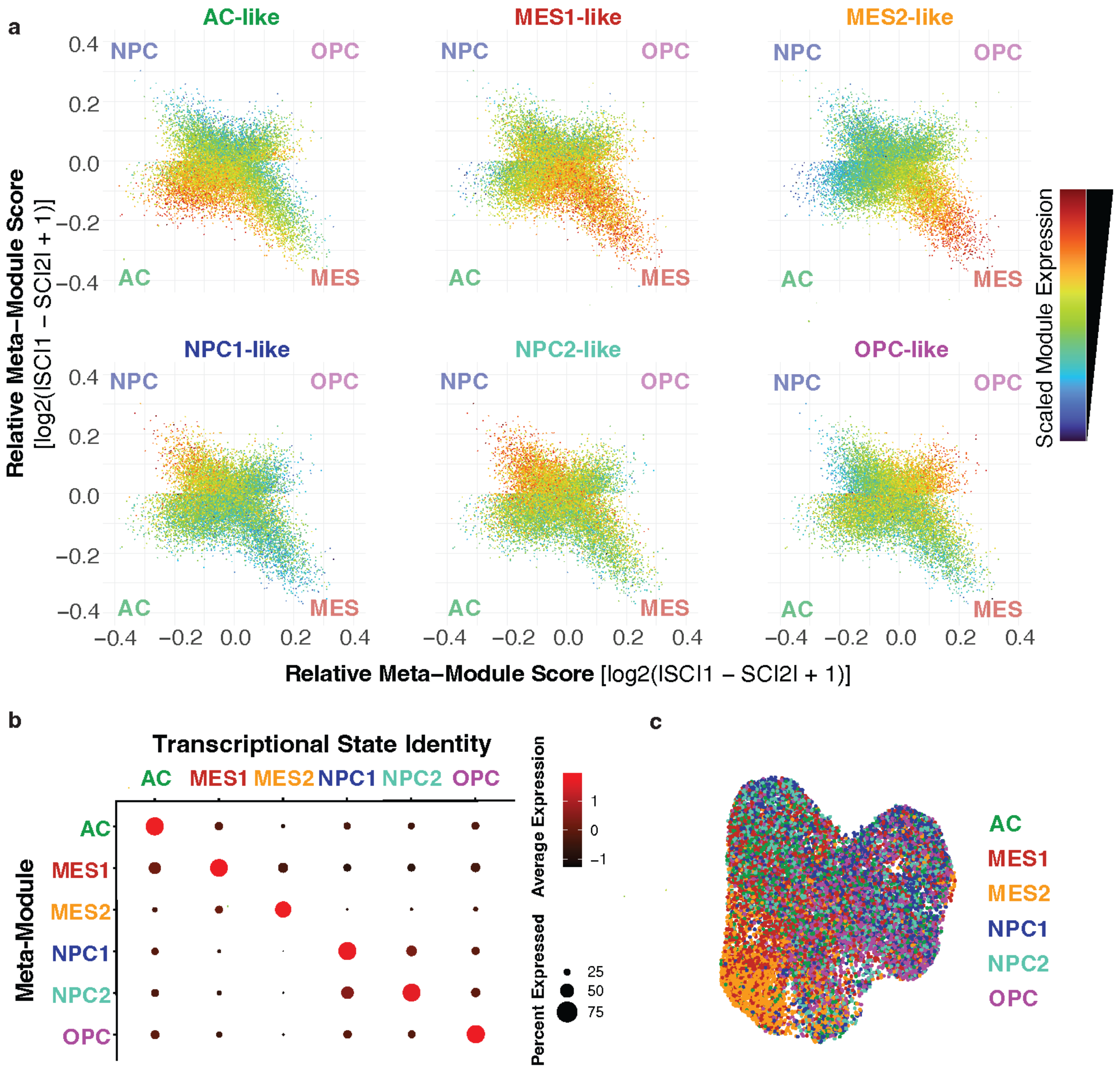
a
AC-like
MES1-like
MES2-like
NPC1-like
NPC2-like
OPC-like
NPC
OPC
AC
MES
Relative Meta−Module Score [log2(lSCl1 − SCl2l + 1)]
Relative Meta−Module Score [log2(lSCl1 − SCl2l + 1)]
Scaled Module Expression
b
Transcriptional State Identity
AC
MES1
MES2
NPC1
NPC2
OPC
Meta-Module
Average Expression
Percent Expressed
c
AC
MES1
MES2
NPC1
NPC2
OPC

**Supplementary Figure S5: GBM22 orthotopic xenografts are transcriptionally heterogeneous. a.** Two-dimensional depiction of relative meta-module scores of GBM22 xenograft cells for the reported GBM cell transcriptional states defined by Neftel et al. (2019). Cells are colored by their enrichment for the six underlying GBM cell transcriptional state meta-modules. Each corner represents a transcriptional state meta-module. **b.** Dot plot of meta-module expression in GBM22 xenograft cells for GBM cell states grouped by assigned transcriptional state identity (6 states) based on predominantly enriched meta-module for each cell. The size of the dot encodes the percentage of cells within each transcriptional state with detected expression of the module, while the color depicts the mean expression level across all cells within each transcriptional state. **c.** UMAP of GBM22 orthotopic xenograft cells colored by assigned transcriptional state identity (6 states).

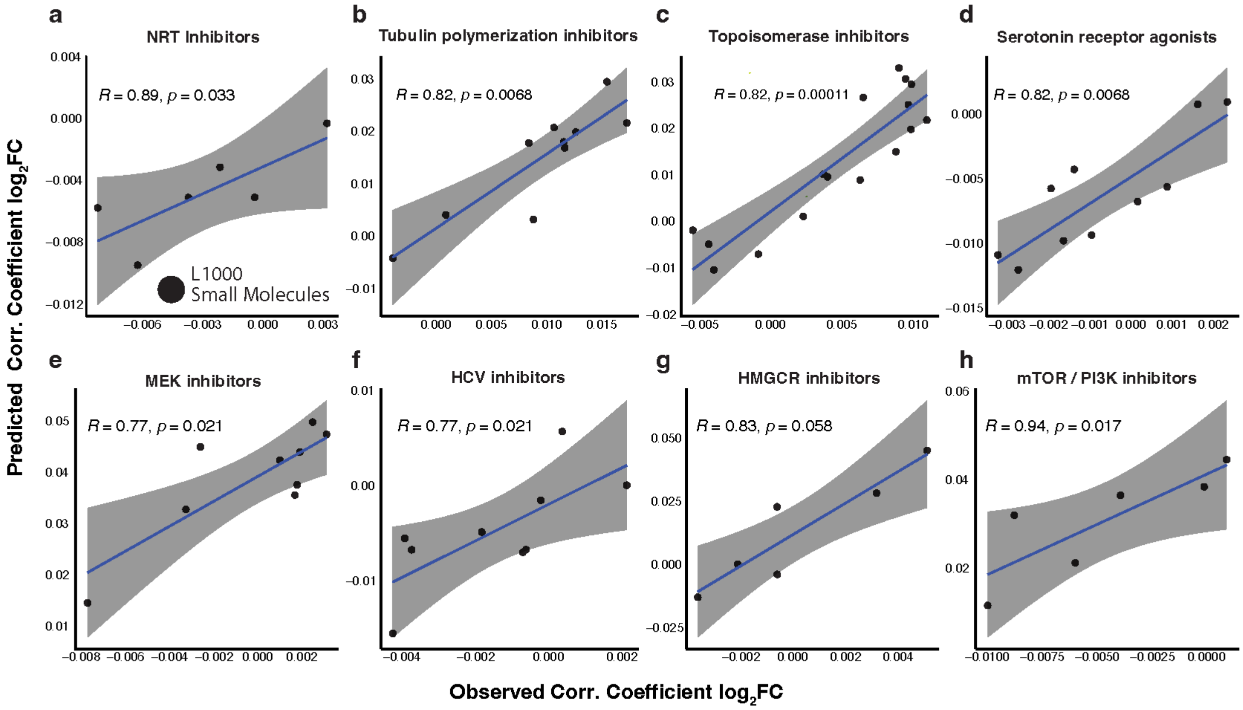


**Supplementary Figure S6: In silico perturbation and predictive differential connectivity analysis models the drug-connectivity perturbation response to alisertib *in vivo*. a-h.** Scatterplots of L1000 small molecule TCSs within individual compound classes depicting ISOSCELES-predicted correlation shift ($log_2FC$) vs. observed correlation shift ($log_2FC$) in alisertib-treated xenografts. Differential small molecule correlations were calculated using *limma*.

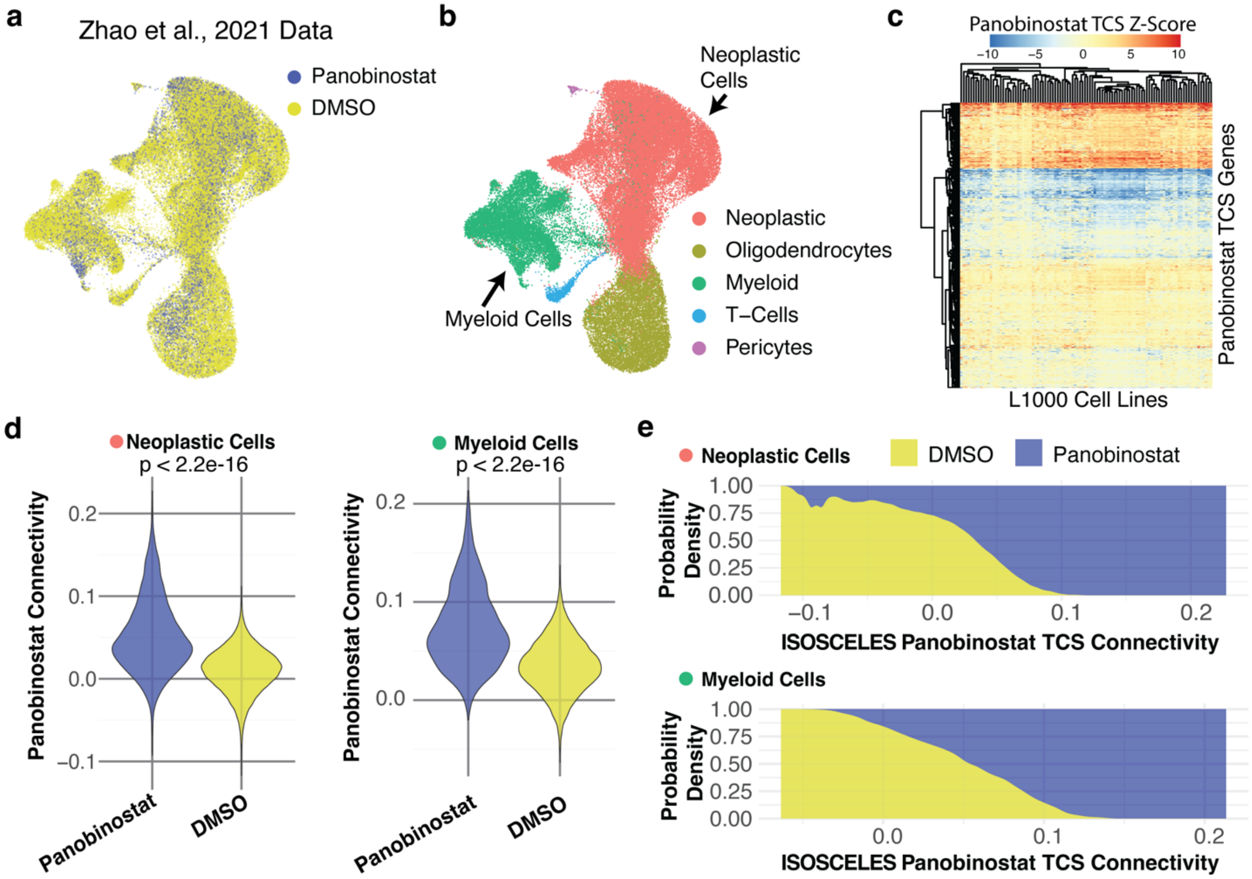


**Supplementary Figure S7: ISOSCELES connectivity analysis with an L1000 derived panobinostat TCS separates vehicle-treated from panobinostat-treated cells in both neoplastic and myeloid cell populations. a.** UMAP of 62,250 single-cell transcriptomes from patient-derived acute slice culture samples obtained from Zhao et al., 2021, treated with DMSO or the HDAC inhibitor panobinostat (0.2 µM). **b.** UMAP of single-cell transcriptomes colored by discrete cell type. **c.** Heatmap showing z-scores of panobinostat-specific gene expression (vs. treatment naïve) for genes retained in the Panobinostat TCS across cell lines tested in the L1000 dataset. **d.** Violin plots showing panobinostat TCS connectivity of Panobinostat-treated vs. DMSO-treated cells in neoplastic and myeloid populations, respectively (Wilcoxon rank sum test with continuity correction, p-value < 2.2e-16). **e.** Area-normalized kernel-density estimate (KDE) plots of ISOSCELES calculated panobinostat TCS connectivity colored by treatment with DMSO or panobinostat, across neoplastic and myeloid cell populations, respectively.

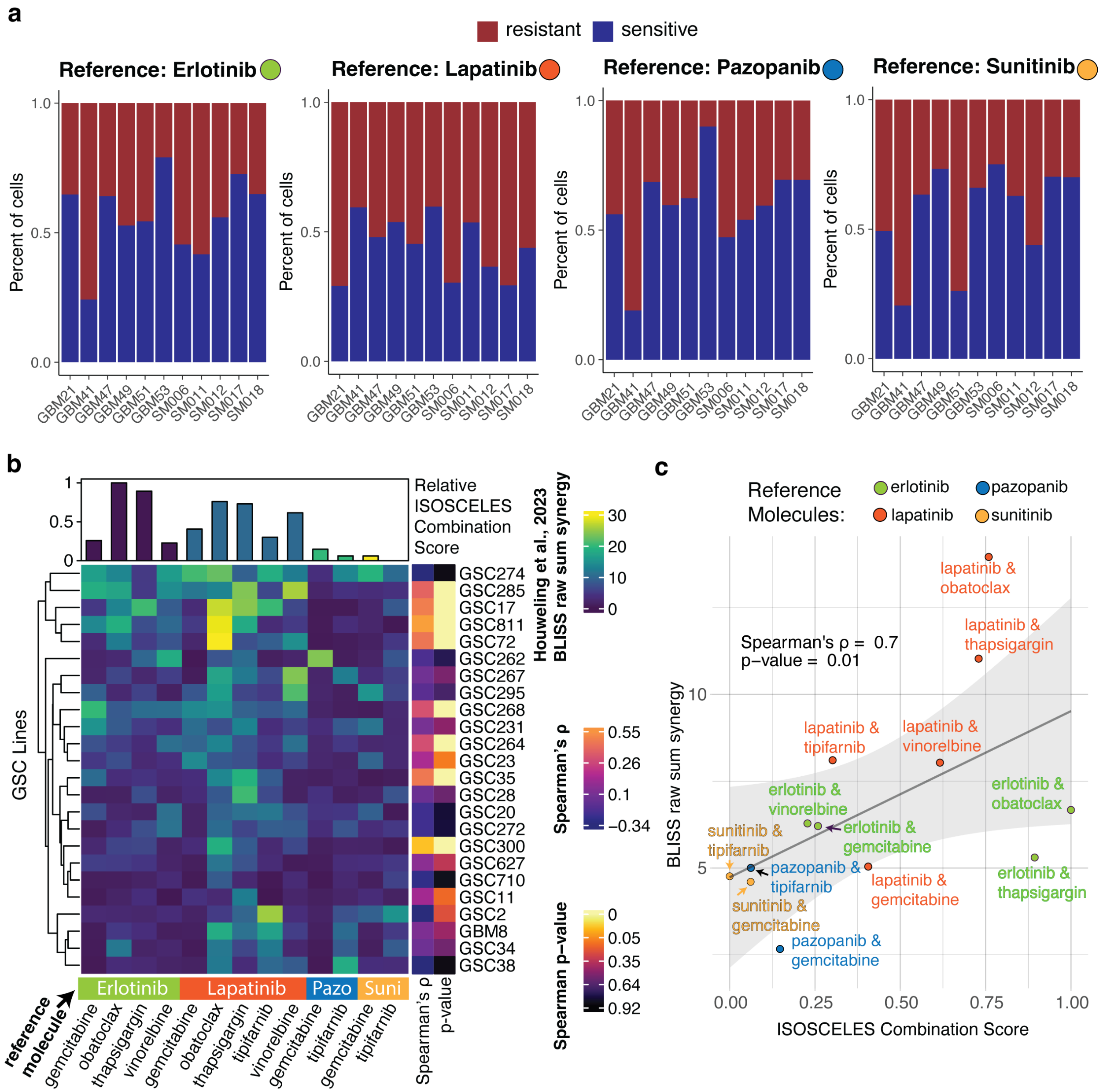
a
resistant
sensitive
Reference: Erlotinib
Reference: Lapatinib
Reference: Pazopanib
Reference: Sunitinib
Percent of cells
1.0
0.5
0.0
GBM21
GBM41
GBM47
GBM49
GBM51
GBM53
SM006
SM011
SM012
SM017
SM018
b
Relative ISOSCELES Combination Score
GSC Lines
GSC274
GSC285
GSC17
GSC811
GSC72
GSC262
GSC267
GSC295
GSC268
GSC231
GSC264
GSC23
GSC35
GSC28
GSC20
GSC272
GSC300
GSC627
GSC710
GSC11
GSC2
GBM8
GSC34
GSC38
Erlotinib
Lapatinib
Pazo
Suni
reference molecule
gemcitabine
obatoclax
thapsigargin
vinorelbine
tipifarnib
Spearman's ρ
p-value
Houweling et al., 2023 BLISS raw sum synergy
30
20
10
0
Spearman's ρ
0.55
0.26
0.1
−0.34
Spearman p−value
0
0.05
0.35
0.64
0.92
c
Reference Molecules:
erlotinib
lapatinib
pazopanib
sunitinib
Spearman's ρ = 0.7
p−value = 0.01
lapatinib & obatoclax
lapatinib & thapsigargin
lapatinib & tipifarnib
lapatinib & vinorelbine
erlotinib & vinorelbine
erlotinib & gemcitabine
erlotinib & obatoclax
sunitinib & tipifarnib
pazopanib & tipifarnib
lapatinib & gemcitabine
erlotinib & thapsigargin
sunitinib & gemcitabine
pazopanib & gemcitabine
BLISS raw sum synergy
ISOSCELES Combination Score
0.00
0.25
0.50
0.75
1.00

**Supplementary Figure S8: ISOSCELES combination scoring predicts relative synergy of small molecule combinations *in vitro*. a.** Stacked bar plots depicting proportion of cells predicted to be sensitive or resistant across individual patient GBM tumors to reference small molecules erlotinib, lapatinib, pazopanib, and sunitinib respectively. **b.** Heatmap of BLISS raw sum synergy of small molecule combinations obtained from Houweling et al., 2023. Top bar plot annotation depicts ISOSCELES calculated combination score using labeled reference molecules depicted on the bottom annotation bar. Sidebar annotations depict Spearman's ρ and p-value between ISOSCELES combination score and observed BLISS raw sum synergy for each individual cell line. **c.** Scatterplot of ISOSCELES calculated combination score and mean BLISS raw sum synergy across all cell lines tested for small molecule combinations screened in Houweling et al., 2023 (Spearman's $\rho = 0.7$, p-value = 0.01). Points are colored by the reference molecule used for combination scoring. Shaded gray area around the line of best fit depicts standard error.

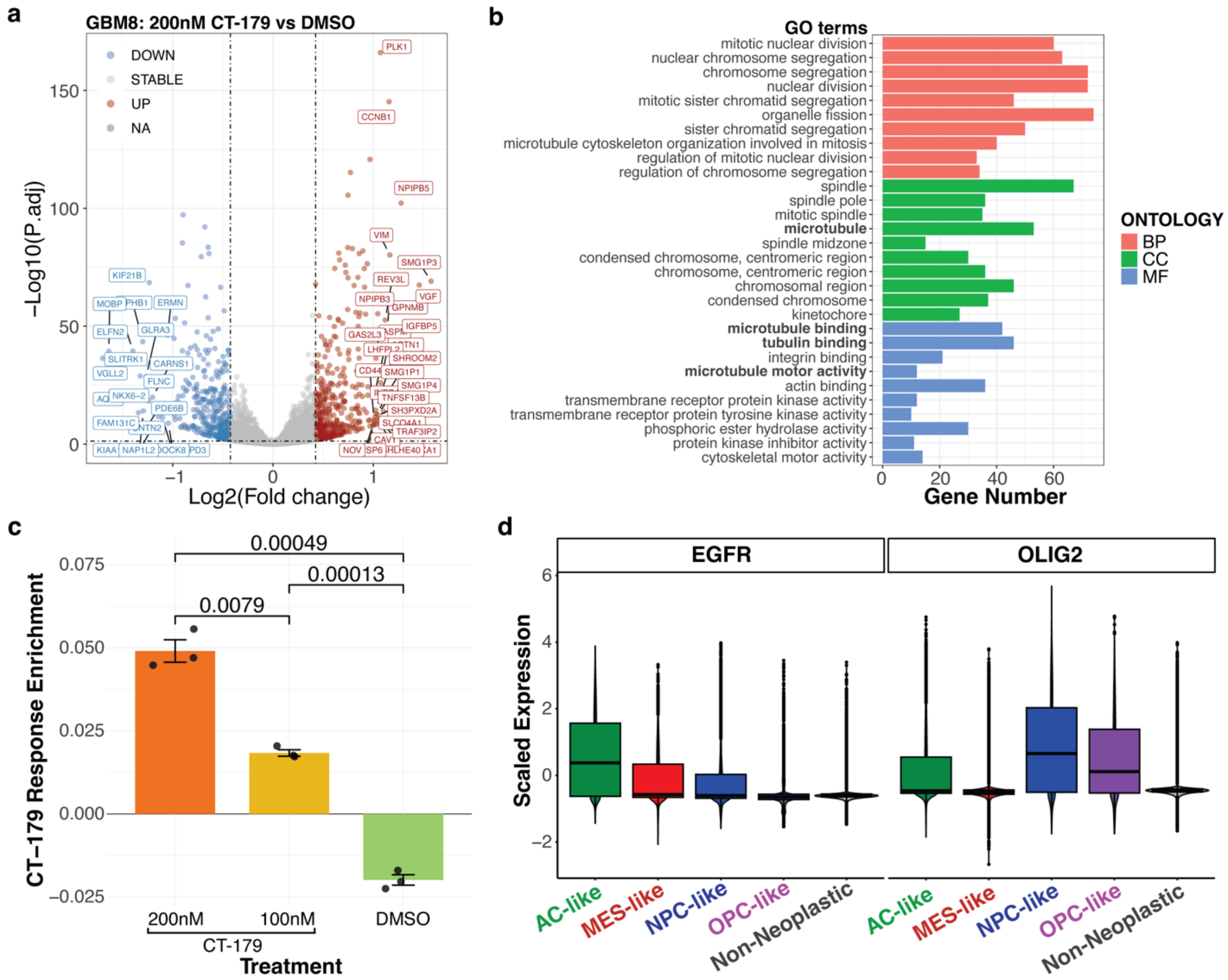

a
GBM8: 200nM CT-179 vs DMSO
DOWN
STABLE
UP
NA
−Log10(P.adj)
Log2(Fold change)
PLK1
CCNB1
NPIPB5
VIM
SMG1P3
REV3L
VGF
NPIPB3
GPNMB
IGFBP5
GAS2L3
SHROOM2
SMG1P1
CD44
SMG1P4
TNFSF13B
SH3PXD2A
TRAF3IP2
NOV
KIF21B
MOBP
PHB1
ERMN
ELFN2
GLRA3
SLITRK1
CARNS1
VGLL2
FLNC
NKX6−2
PDE6B
FAM131C
NAP1L2
b
GO terms
mitotic nuclear division
nuclear chromosome segregation
chromosome segregation
nuclear division
mitotic sister chromatid segregation
organelle fission
sister chromatid segregation
microtubule cytoskeleton organization involved in mitosis
regulation of mitotic nuclear division
regulation of chromosome segregation
spindle
spindle pole
mitotic spindle
microtubule
spindle midzone
condensed chromosome, centromeric region
chromosome, centromeric region
chromosomal region
condensed chromosome
kinetochore
microtubule binding
tubulin binding
integrin binding
microtubule motor activity
actin binding
transmembrane receptor protein kinase activity
transmembrane receptor protein tyrosine kinase activity
phosphoric ester hydrolase activity
protein kinase inhibitor activity
cytoskeletal motor activity
Gene Number
ONTOLOGY
BP
CC
MF
c
CT−179 Response Enrichment
0.00049
0.00013
0.0079
200nM
100nM
DMSO
CT-179
Treatment
d
EGFR
OLIG2
Scaled Expression
AC-like
MES-like
NPC-like
OPC-like
Non-Neoplastic

**Supplementary Figure S9: Generation of a GBM CT-179 transcriptional response signature.** **a.** Volcano plot of differential expression between 200nM CT-179 and vehicle treated GBM8 cells calculated via DESeq2. **b.** Barplot depicting gene ontology analysis of the filtered CT-179 transcriptional response signature. Bars are colored by the respective ontology used. Selected GO terms highlighted include 'microtubule', 'microtubule binding', 'tubulin binding', and 'microtubule motor activity'. **c.** Barplot of singscore-calculated enrichment for the directional CT-179 transcriptional response signature in 200nM CT-179 treated, 100nM CT-179 treated, and vehicle-treated GBM8 cells after 24 hours. Significance bars depict p-value following pairwise t-tests (n = 3 per treatment group). **d.** Box and violin plots depicting the expression of EGFR and OLIG2 in patient GBM tumor cells grouped by assigned Neftel et al. GBM cell transcriptional state.
